# Nuclear retention and RNA editing safeguard self-tolerance to immunogenic intron lariats

**DOI:** 10.64898/2026.11.08.765213

**Authors:** Keonyong Lee, Jayoung Ku, Soo Young Park, Tria Asri Widowati, Moonhyeon Jeon, Namwook Kim, Hyukjun Chung, Min Ji Kim, Da Sol Kim, Ha Yeon Kim, Hyewon Chung, Woo Dae Jang, Han Suk Ryu, Yoosik Kim

**Author notes:** Correspondence: Han Suk Ryu; Yoosik Kim. These authors contributed equally (Keonyong Lee, Jayoung Ku, Soo Young Park).

## Abstract

Introns are excised as lariats and rapidly degraded upon linearization, leaving their post-splicing role unclear. Here, we investigate their immunoregulatory potential by analyzing the intron lariat landscape upon inhibition of the debranching enzyme DBR1. We find that lariats harboring transposable elements (TEs), especially inverted *Alu* repeats (IR*Alus*), elicit cell cycle-dependent activation of protein kinase R (PKR), but only in cells harboring a high burden, such as breast cancer. These immunogenic lariats are initially retained in the nucleus, but upon cytosolic release, they drive PKR activation and apoptosis. Exploiting this, we combine DBR1 depletion with anti-mitotic chemotherapy to enhance lariat-PKR interactions and promote cancer cell death in vitro and in vivo. We further show that A-to-I editing of IR*Alus*-containing lariats weakens their interaction with PKR and suppresses PKR activation. Our findings highlight two parallel molecular mechanisms to sustain self-tolerance to repeat-rich introns, revealing a vulnerability exploitable for anti-tumor therapy.

## INTRODUCTION

Introns represent a distinguishing feature of eukaryotes.^1,2^ Introns vary significantly in number and size across species, with 95% of human genes having eight introns on average.^3–5^ During splicing, introns are released as a lariat and undergo debranching by debranching RNA lariats 1 (DBR1), followed by exonuclease-mediated degradation.^6–8^ One potential reason behind the rapid degradation is that intron RNAs may act as cellular double-stranded RNAs (dsRNAs) by hosting transposable elements (TEs), including short interspersed nuclear elements (SINEs) and long interspersed nuclear elements (LINEs). Growing evidence suggests that TE-derived RNAs can form dsRNAs recognized as non-self by pattern recognition receptors, including protein kinase R (PKR) and melanoma differentiation-associated protein 5 (MDA5), and trigger antiviral signaling.^9–13^ More importantly, 60% of *Alu*, a primate-specific SINE with over one million copies in the human genome, are located in intronic regions, and inverted *Alu* repeats (IR*Alus*) constitute the most abundant class of cellular dsRNAs, suggesting that intron RNAs may pose a major immunoregulatory challenge to cells.^14,15^

To prevent aberrant immunogenic effects, intron-derived cellular dsRNAs are tightly regulated through multiple mechanisms. Upon inhibition of nuclear RNA decay by phosphorothioate DNAs, intronic and intergenic repeats of TEs accumulate and activate dsRNA sensors.^16^ Importantly, numerous studies exploited the immunogenic potential of intronic RNAs for cancer therapy. For example, epigenetic therapy, including the DNA demethylating agent, 5-aza-2′-deoxycytidine, reactivates transcription of intronic TEs, resulting in anti-tumor responses through viral mimicry,^17–20^ and splicing disturbances similarly trigger dsRNA-induced innate immune responses by accumulating intron-retained mRNAs.^21–23^ On the contrary, some studies suggest that post-spliced intron RNAs may exert immunosuppressive effects. For example, depletion or mutation of the *DBR1* gene confers susceptibility to viral infection by impairing stress granule assembly and/or elevating the expression of imperfect IR*Alus* dsRNAs with weaker immunogenicity than perfect dsRNAs, subsequently inhibiting PKR.^24–27^ Yet, molecular details of intron lariats, such as their TE landscape, localization, editing, and potential direct interaction with RNA sensors, remain largely unexplored.

Here, we investigated the effects of intron lariat accumulation upon DBR1 depletion across multiple cell lines, focusing on embedded TEs. We found that a substantial increase in TE-harboring introns is required to elicit a strong immunogenic response. Through a comprehensive analysis of the PKR-intron lariat interaction landscape, we found that the type of embedded TEs and subcellular localization govern PKR binding, which we used to uncover cell cycle-dependent PKR activation. Lastly, we analyzed how A-to-I editing of intron RNAs modulates PKR sensing and downstream immune activation. Collectively, our work establishes multi-layered safeguards, including rapid turnover, localization, and editing, against self-intron RNA immunogenicity and proposes DBR1 as a promising target to enhance anti-mitotic cancer therapies.

## RESULTS

### DBR1 inhibition elevates intronic TE-derived dsRNAs

Considering that debranching is the rate-limiting step in intron degradation,^28,29^ we investigated the effect of DBR1 depletion on the expression and recognition of excised intron lariats. We used MDA-MB-231 human triple-negative breast cancer (TNBC) cells as our experimental system, following a previous report showing that spliceosome inhibition results in viral mimicry of intron-retained mRNAs in TNBC.^23^

Following DBR1 depletion using RNAi, selected intron RNAs that were reported to be upregulated in human DBR1 mutant fibroblasts^24^ accumulated as lariat structures, confirmed by divergent reverse transcription-qPCR and RNase R treatment, which selectively degrades linear RNAs **(Extended Data Fig. 1a-d)**. We then performed total RNA-seq over varying durations of DBR1 depletion and found that overall levels of intron RNAs increased over time **(Fig. 1a and Extended Data Fig. 1e)**. Our analysis revealed that intronic RNAs were more enriched upon RNase R treatment in DBR1-deficient cells than in control cells, indicating that DBR1 inhibition globally elevates the levels of intron RNAs in the lariat form **(Fig. 1b)**. In addition, we found that all TEs were significantly upregulated in DBR1-deficient cells **(Fig. 1c)**. Notably, only the intron-embedded TEs were significantly upregulated, whereas TEs in non-intronic regions were unaffected by DBR1 depletion **(Fig. 1d and Extended Data Fig. 1f-i)**. As examples, sequencing reads mapped to the *PTBP1* intron 13 (containing multiple *Alu*s) and *RDH11* intron 4 (containing a LINE1 (L1)) are shown in **Extended Data Fig. 1h**.

**Fig. 1.**
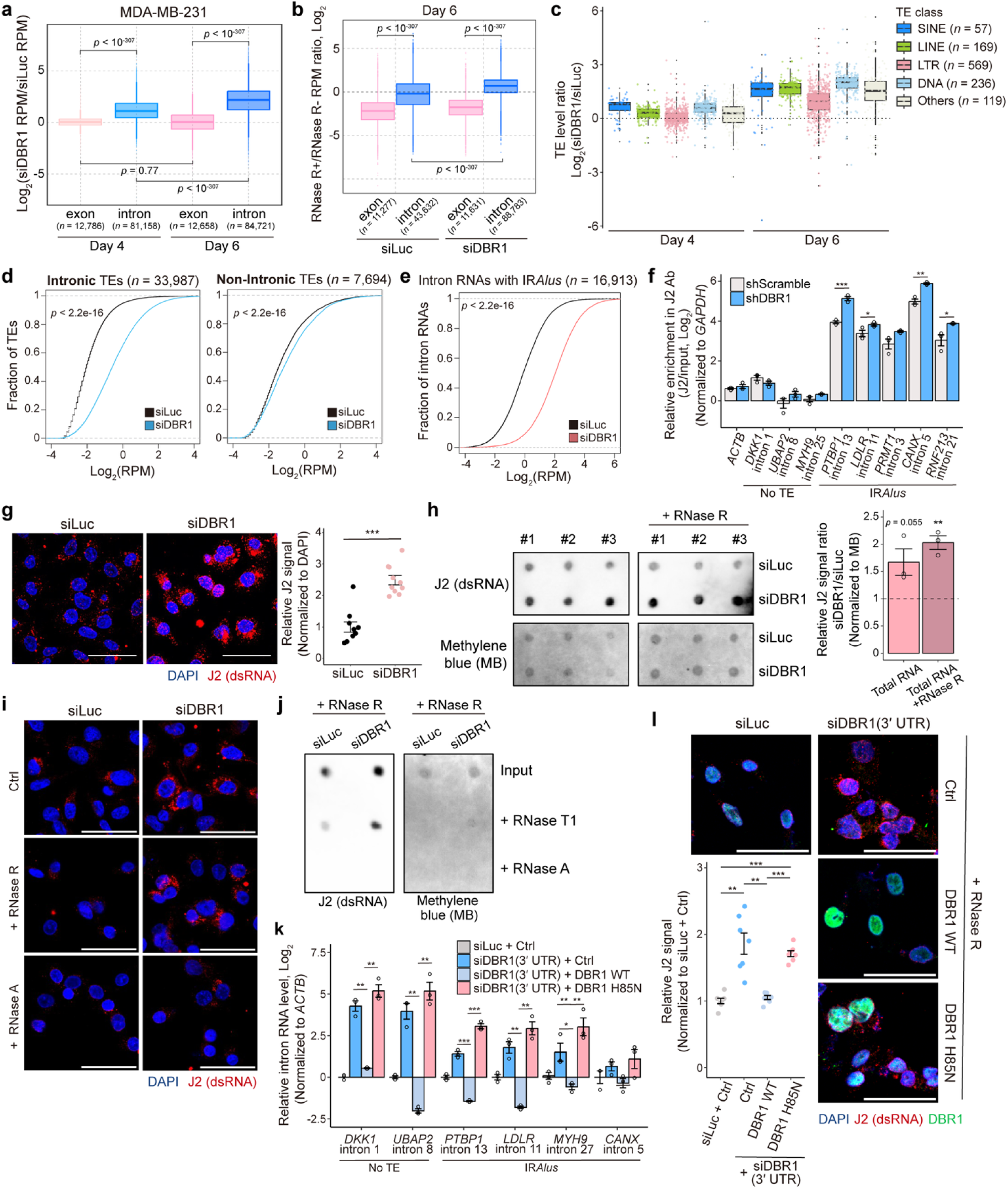
Accumulation of intronic TE-derived dsRNAs upon DBR1 depletion. **a.** Fold change in mean reads per million (RPM) of exonic or intronic regions from total RNA-seq, 4 and 6 days after siDBR1 transfection to MDA-MB-231. **b.** Exon and intron RPM ratios (RNase R-treated vs. untreated total RNA-seq), 6 days after siRNA transfection to MDA-MB-231. **c.** TE expression changes 4 and 6 days after siDBR1 transfection to MDA-MB-231. Each point denotes a specific TE sub-family. **d, e.** Cumulative distribution of mean RPMs for intronic TEs (**d**, left, *n* = 33,987), non-intronic TEs (**d**, right, *n* = 7,694), and IR*Alus*-containing intron RNAs (**e**, *n* = 16,913) in MDA-MB-231 transfected with siRNAs for 6 days. **f.** RT-qPCR analysis of intron RNAs pulled down with J2 from shRNA-transduced MDA-MB-231, normalized to input (*n* = 3). **g.** J2 Immunocytochemistry of MDA-MB-231, 6 days after siRNA transfection. Quantification is shown on the right (*n* = 10). **h.** J2 dot-blot assay of total and RNase R-treated RNAs from MDA-MB-231 transfected with siRNAs for 6 days. Three biological replicates are shown with quantified data on the right and methylene blue staining as the loading control. **i.** J2 signal after RNase R or RNase A treatment in siRNA-transfected MDA-MB-231. **j.** J2 dot-blot of RNase R-treated RNAs from MDA-MB-231, 6 days after siRNA transfection, followed by additional RNase T1 or RNase A treatment. **k.** RT-qPCR of selected intron RNAs following overexpression of DBR1 WT or H85N mutant in DBR1-deficient MDA-MB-231 (*n* = 3). **l.** J2 signal (RNase R-treated) in DBR1-deficient MDA-MB-231 overexpressing DBR1 WT or H85N. Quantification is shown on the left (*n* = 7). All error bars denote s.e.m. All scale bars indicate 50 μm. *P*-values were calculated using either the two-sided Wilcoxon signed-rank test (a, b), the two-sided Mann-Whitney test (d, e), or the one-sided Student’s t-test (f, g, h, k, l); ^∗^*p* < 0.05, ^∗∗^*p* < 0.01, ^∗∗∗^*p* < 0.001.

Considering that IR*Alus* are the primary source of endogenous dsRNAs in humans and are highly abundant in introns, we examined whether DBR1 depletion affects cellular dsRNA expression. Indeed, IR*Alus*-containing intron RNAs increased substantially in our sequencing data **(Fig. 1e and Extended Data Fig. 1j)**. Additionally, IR*Alus*-containing intron RNAs were significantly enriched in the dsRNA-specific J2 antibody pull-down compared to those lacking TEs, and this enrichment was further elevated upon DBR1 depletion **(Fig. 1f)**. Notably, while DBR1 depletion increased total RNA levels **(Extended Data Fig. 1k),** the magnitude was much greater for dsRNAs **(Fig. 1g, h and Extended Data Fig. 1l)**. The RNase R treatment only had a marginal effect on reducing the dsRNA signal in DBR1-deficient cells **(Fig. 1h, i)**. Even total RNA levels remained higher in DBR1-deficient cells after RNase R treatment, indicating that both increased dsRNAs and total RNAs are derived from intron lariats **(Extended Data Fig. 1k)**. The RNase A treatment at high concentration, which degrades both ssRNAs and dsRNAs, abolished both the dsRNA and total RNA signals **(Fig. 1i and Extended Data Fig. 1k)**. To further verify increased dsRNA expression upon DBR1 depletion, we examined the nuclease sensitivity. Equal amounts of RNase R-treated total RNA from siLuc- and siDBR1-transfected cells were treated with RNase T1 (degrades ssRNAs) or RNase A, and equal volumes of each reaction were analyzed by J2 dot-blot assay. Whereas the remaining RNA in siLuc samples was largely degraded by both nucleases, siDBR1 samples retained a J2 signal that was resistant to RNase T1 but sensitive to RNase A, indicating that the RNase R-resistant RNAs accumulating upon DBR1 depletion exist substantially in a dsRNA conformation **(Fig. 1j)**.

To confirm the DBR1-dependency for the observed dsRNA accumulation, we ectopically overexpressed WT (DBR1 WT) and catalytic mutant DBR1 (DBR1 H85N),^24^ respectively, in DBR1-deficient cells **(Extended Data Fig. 1m)**. While the introduction of DBR1 WT rescued intron RNA and dsRNA levels, DBR1 H85N did not **(Fig. 1k, l)**. Moreover, DBR1 depletion did not affect the level of mitochondrial RNAs (mtRNAs), another major source of endogenous dsRNAs, ruling out mtRNAs as a contributing source of the increased dsRNAs **(Extended Data Fig. 1n)**.

### Intron-mediated PKR activation triggers apoptosis upon DBR1 depletion

Next, we examined the downstream consequence of intron lariat RNA accumulation. We found that DBR1 depletion triggered cell death and increased the levels of apoptotic markers, including cleaved PARP (c-PARP), cleaved Caspase-8 (c-Caspase-8), and cleaved Caspase-3 (c-Caspase-3), in MDA-MB-231 **(Fig. 2a, b and Extended Data Fig. 2a, b)**. To identify the responsible signaling pathway, we individually downregulated dsRNA-associated proteins (PKR, MDA5, RIG-I, RNase L, NF90/NF110, PACT, and TRBP) in DBR1-deficient cells **(Extended Data Fig. 2c, d)**. Interestingly, PKR and NF90/NF110 depletion significantly mitigated cell death in DBR1-deficient cells **(Fig. 2c)**. Given the well-established role of PKR in dsRNA-mediated cell death,^30^ we focused on its involvement and found that DBR1 downregulation had a much milder effect on cell viability in MDA-MB-231 PKR KO cells **(Fig. 2d and Extended Data Fig. 2e)**.

**Fig. 2.**
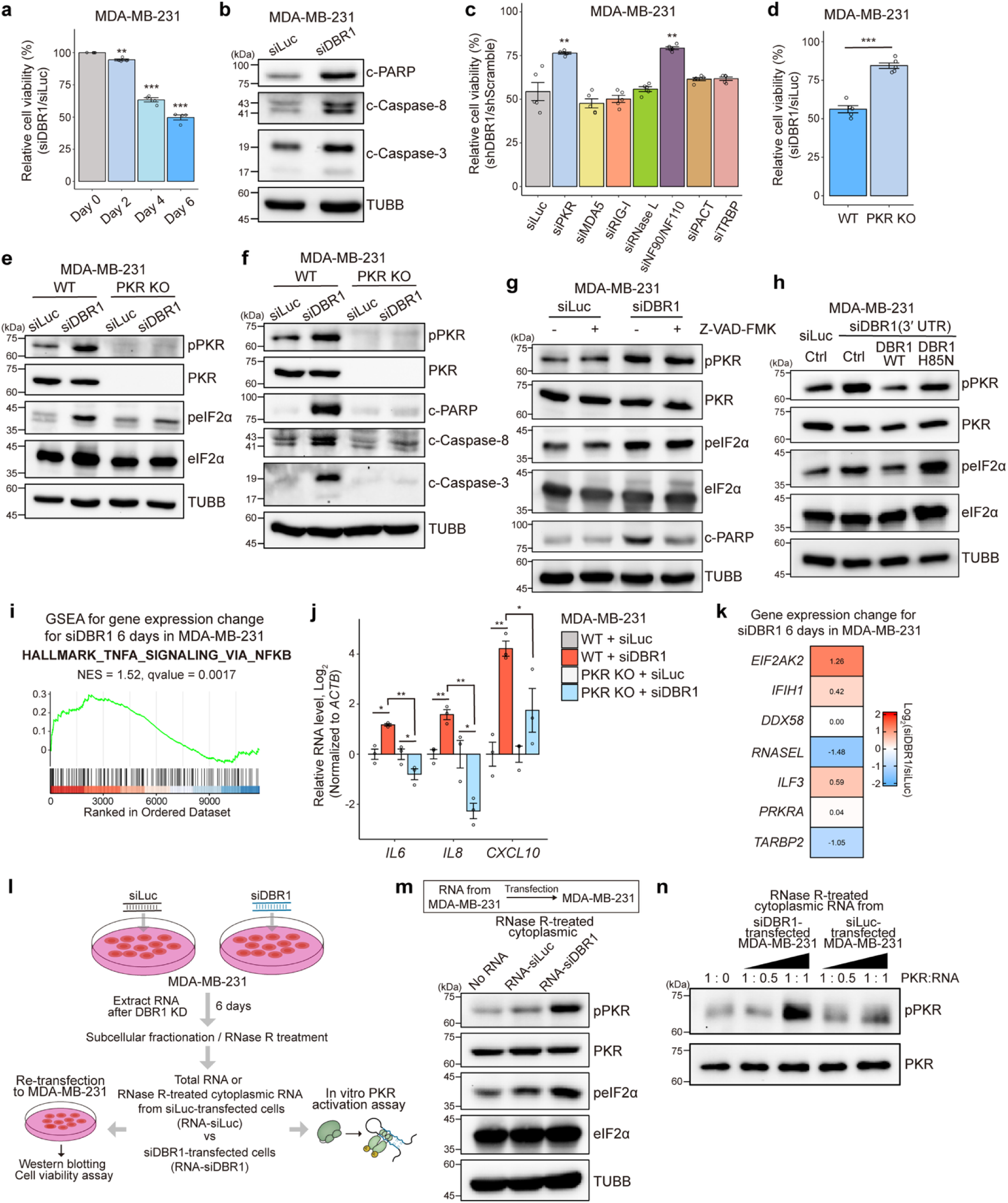
DBR1 depletion-induced cell death via PKR. **a.** Cell viability of MDA-MB-231 up to 6 days after siDBR1 transfection (*n* = 4). **b.** Western blotting (WB) of apoptotic markers (c-PARP, c-Caspase-8, and c-Caspase-3) 4 days after siDBR1 transfection to MDA-MB-231. **c.** Cell viability upon knockdown of dsRNA-associated proteins in DBR1-deficient MDA-MB-231 (*n* = 5). **d.** Cell viability 4 days after siDBR1 transfection to MDA-MB-231 WT and PKR KO (*n* = 5). **e, f.** WB of PKR and eIF2α phosphorylation (**e**) and apoptotic markers (**f**) 4 days after siDBR1 transfection to MDA-MB-231 WT and PKR KO. **g.** Effect of Z-VAD-FMK treatment on PKR and eIF2α phosphorylation and PARP cleavage in DBR1-deficient MDA-MB-231. **h.** Effect of DBR1 WT or H85N overexpression on PKR and eIF2α phosphorylation in DBR1- deficient MDA-MB-231. **i.** GSEA results for HALLMARK_TNFA_SIGNALING_VIA_NFKB gene set from 6 days siDBR1-transfected MDA-MB-231. **j.** RT-qPCR of *IL6, IL8,* and *CXCL10* mRNAs 6 days after siDBR1 transfection to MDA-MB-231 WT and PKR KO (*n* = 3). **k.** Heatmap of dsRNA-associated gene expression from RNA-seq analysis of MDA-MB-231 transfected with siDBR1 for 6 days. **l.** Experimental scheme of RNA re-transfection and in vitro PKR activation assay using total RNA or RNase R-treated cytoplasmic RNA extracted from MDA-MB-231 transfected with siRNAs for 6 days (RNA-siDBR1 or RNA-siLuc). **m.** WB of PKR and eIF2α phosphorylation 24 h after re-transfection of RNase R-treated cytoplasmic RNA-siDBR1 or RNA-siLuc into MDA-MB-231. **n.** In vitro PKR activation assay using recombinant PKR and RNase R-treated cytoplasmic RNA-siDBR1 or RNA-siLuc. The ratio indicates the mass ratio of RNA to PKR. All error bars denote s.e.m. *P*-values were calculated using the one-sided Student’s t-test (a, c, d, j); ^∗^*p* < 0.05, ^∗∗^*p* < 0.01, ^∗∗∗^*p* < 0.001.

Upon binding to dsRNAs, PKR undergoes activation via autophosphorylation, and phosphorylated PKR (pPKR) phosphorylates the α subunit of eukaryotic initiation factor 2 (eIF2α) and triggers apoptosis by activating the caspase-8/caspase-3 pathway.^31,32^ We found that the loss of DBR1 increased the pPKR and phosphorylated eIF2α (peIF2α) **(Fig. 2e and Extended Data Fig. 2f, g)**. The induction of apoptotic markers and peIF2α were significantly reduced in PKR KO cells, further confirming that PKR mediates DBR1 depletion-induced apoptosis **(Fig. 2e, f and Extended Data Fig. 2h)**. To exclude the possibility that PKR activation is an indirect consequence of apoptosis, we treated cells with the pan-caspase inhibitor, Z-VAD-FMK, which rescued cell death and PARP cleavage without affecting pPKR and peIF2α **(Fig. 2g and Extended Data Fig. 2i)**. Moreover, introducing DBR1 WT fully rescued PKR and eIF2α phosphorylation, whereas the DBR1 H85N did not, suggesting that inhibiting the intron debranching process activates PKR signaling **(Fig. 2h)**.

To further characterize the transcriptome-wide effects of DBR1 depletion, we analyzed gene expression profiles in DBR1-deficient MDA-MB-231. Gene set enrichment analysis (GSEA) revealed significant upregulation of tumor necrosis factor α (TNFα) signaling via the nuclear factor κB (NF-κB) pathway, a downstream target of PKR **(Fig. 2i and Extended Data Fig. 2j)**.^33^ We experimentally confirmed the upregulation of NF-κB-responsive genes, including *IL6*, *IL8*, and *CXCL10*, in DBR1-deficient MDA-MB-231 cells **(Fig. 2j)**. Moreover, the DBR1 depletion-mediated induction of NF-κB-responsive genes was significantly reduced in MDA-MB-231 PKR KO cells **(Fig. 2j)**. Interestingly, among the dsRNA-associated genes examined above in **Fig. 2c**, *EIF2AK2* showed the greatest upregulation upon DBR1 depletion, further highlighting the involvement of PKR in DBR1 depletion-induced responses **(Fig. 2k)**. Overall intron RNA levels were comparable between MDA-MB-231 WT and PKR KO cells after DBR1 depletion, indicating that the attenuated apoptosis and inflammatory responses in PKR KO cells were not due to differential accumulation of intron RNAs **(Extended Data Fig. 2k)**. Alongside TNFα and NF-κB-mediated inflammatory responses, DBR1-deficient MDA-MB-231 cells showed enrichment of cell cycle-related responses, particularly the G2/M checkpoint responses, while showing no enrichment in splicing-altered genes **(Extended Data Fig. 2j)**.

We then investigated whether intron lariat RNAs can directly interact and activate PKR. First, we performed an RNA re-transfection assay using total RNAs extracted from control and DBR1-knockdown cells **(Fig. 2l)**. Transfection of RNAs extracted from siDBR1-transfected cells (Total RNA-siDBR1) successfully increased overall intron RNA levels in recipient cells, whereas control RNAs (Total RNA-siLuc) did not **(Extended Data Fig. 2l)**. Notably, transfection of total RNA-siDBR1 triggered stronger PKR and eIF2α phosphorylation and induced cell death compared to total RNA-siLuc **(Extended Data Fig. 2m, n)**. Next, an in vitro PKR activation assay using recombinant PKR revealed that incubation with total RNA-siDBR1 phosphorylated PKR in vitro more effectively than total RNA-siLuc **(Extended Data Fig. 2o)**. To further confirm whether these PKR-activating RNAs are in the cytoplasm, we isolated cytoplasmic RNA fractions and treated them with RNase R **(Fig. 2l and Extended Data Fig. 2p, q)**. Consistent with the total RNA experiments, RNase R-treated cytoplasmic RNA-siDBR1 activated PKR in cells and in vitro more potently than cytoplasmic RNA-siLuc **(Fig. 2m, n)**.

### Sufficient TE-harboring intron RNA accumulation may trigger PKR activation

While our data clearly show that DBR1 depletion results in PKR-dependent cell death in MDA-MB-231, recent studies suggest an immunosuppressive effect of intron lariats, reporting that inborn errors of *DBR1* confer susceptibility to viral infection.^24,25^ Mechanistically, DBR1 depletion disrupts stress granule formation, thereby impairing PKR activation upon infection.^26^ To resolve this discrepancy, we downregulated DBR1 in HEK-293T, the experimental system used in the previous study, and found that DBR1 depletion did not trigger PKR and eIF2α phosphorylation **(Fig. 3a).** We next examined intron lariat-derived dsRNA levels in DBR1-deficient HEK-293T and MDA-MB-231 using immunocytochemistry following RNase R treatment. While DBR1 depletion markedly increased dsRNA levels in MDA-MB-231, a moderate increase was observed in HEK-293T **(Fig. 3b)**. Consistently, RT-qPCR analysis revealed less accumulation of intron RNAs in HEK-293T **(Extended Data Fig. 3a)**. Moreover, RNase R-treated cytoplasmic RNAs from DBR1-deficient HEK-293T failed to activate PKR in vitro and when transfected into MDA-MB-231 **(Fig. 3c, d)**.

**Fig. 3.**
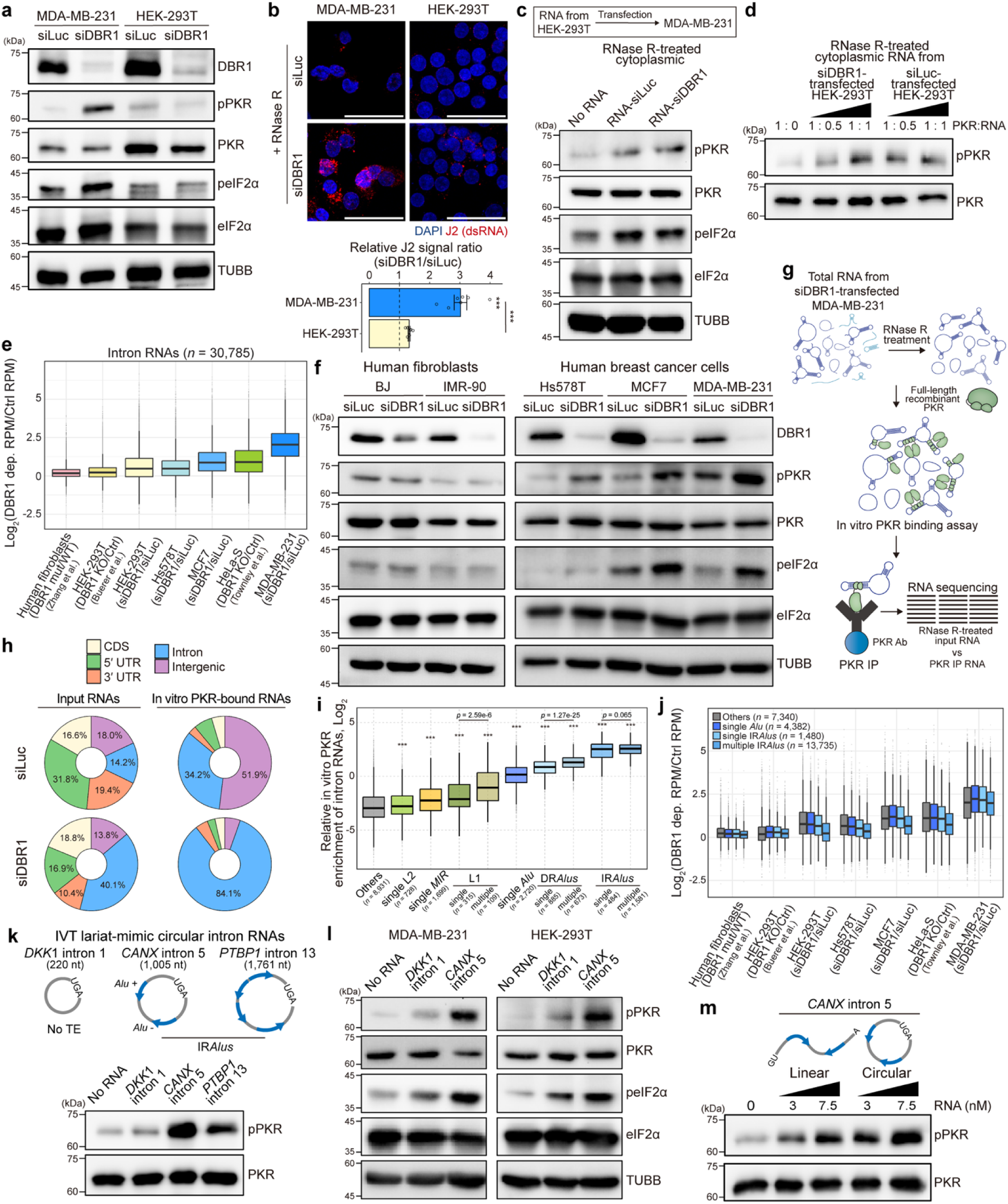
Differential intron RNA accumulation across cell types and characterization of PKR- interacting lariat RNAs. **a.** WB of PKR and eIF2α phosphorylation 6 days after siDBR1 transfection to MDA-MB-231 and HEK-293T. **b.** J2 signal (RNase R-treated) 6 days after siRNA transfection to MDA-MB-231 and HEK-293T. Quantification is shown on the right (*n* = 7). Scale bar indicates 50 μm. **c.** WB of PKR and eIF2α phosphorylation 24 h after re-transfection of RNase R-treated cytoplasmic RNA-siDBR1 or RNA-siLuc (extracted from HEK-293T) to MDA-MB-231. **d.** In vitro PKR activation assay using recombinant PKR and RNase R-treated cytoplasmic RNA-siDBR1 or RNA-siLuc extracted from HEK-293T. The ratio indicates the mass ratio of RNA to PKR. **e.** Box plot of intron RNA accumulation across different cell types upon DBR1 depletion or mutation, calculated by Log_2_(RPM in DBR1-depleted or mutant / control cells) for 30,785 intron RNAs. For siDBR1-transfected cell lines (HEK-293T, Hs578T, MCF7, and MDA-MB-231), RNA-seq data were collected 6 days after transfection. RNA-seq data for DBR1-mutant fibroblasts, DBR1 KO HEK-293T, and DBR1 KO HeLa-S were obtained from published studies^24,34,35^. **f.** Comparison of PKR and eIF2α phosphorylation 6 days after siDBR1 transfection to human fibroblasts (BJ and IMR-90) and breast cancer cells (Hs578T, MCF7, and MDA-MB-231), confirmed by WB. **g.** Schematic representation of an in vitro PKR binding assay followed by RNA-seq (in vitro PKR binding seq). 5% of RNase R-treated input RNA was used as a control. **h.** Proportions of genomic region-mapped reads in the in vitro PKR binding-seq for RNase R-treated total RNA extracted from siRNA-transfected MDA-MB-231. **i.** Relative in vitro PKR enrichment of intron RNAs (in vitro PKR RPM/input RPM) from siDBR1- transfected sample, depending on the type and the number of TEs (*MIR*, *Alu*, L1, L2, DR*Alus*, and IR*Alus*). “Others” indicate RNAs that lack *MIR*, *Alu*, L1, and L2. **j.** Box plot of the intron RNA fold change across different cell types upon DBR1 depletion/ mutation, calculated by Log_2_(RPM in DBR1-depleted or mutant / control cells) and grouped by the presence and number of IR*Alus*. **k.** In vitro PKR activation assay using recombinant PKR and in vitro synthesized lariat-mimic circular intron RNAs (7.5 nM): *DKK1* intron 1 (220 nt, no TE), *CANX* intron 5 (1,005 nt, with IR*Alus*), and *PTBP1* intron 13 (1,761 nt, with IR*Alus*). **l.** WB of PKR and eIF2α phosphorylation in MDA-MB-231 and HEK-293T transfected with circular *DKK1* intron 1 or *CANX* intron 5 RNA (250 ng/mL), assessed at 12 h post-transfection. **m.** In vitro PKR activation assay using recombinant PKR and in vitro synthesized linear or circular *CANX* intron 5 RNA. All error bars denote s.e.m. *P*-values were calculated using either the one-sided Student’s t-test (b) or the two-sided Wilcoxon signed-rank test (i); ^∗^*p* < 0.05, ^∗∗^*p* < 0.01, ^∗∗∗^*p* < 0.001.

We then compared RNA-seq data from published studies^24,34,35^ and our own and found that the extent of intron RNA accumulation varied across cell lines **(Fig. 3e)**. Particularly, DBR1 mutant patient-derived fibroblasts and DBR1-deficient HEK-293T have a significantly lower degree of intron RNA accumulation compared to those of cancer cell lines **(Fig. 3e)**. In addition, overall TE expression was not significantly increased in DBR1-deficient HEK-293T **(Extended Data Fig. 3b)**. These data indicate that a sufficient increase in intron lariats, especially those with TEs, might be required to activate PKR. Consistent with our model, when we compared the effect of DBR1 depletion in two human fibroblast cell lines (BJ and IMR-90) and three breast cancer cell lines (Hs578T, MCF7, and MDA-MB-231), the extent of PKR activation was positively correlated with the degree of intron RNA and TE accumulation **(Fig. 3f and Extended Data Fig. 3b)**. The accumulation of intron lariat-derived dsRNAs was more pronounced in MDA-MB-231 and MCF7, which also exhibited robust PKR activation **(Fig. 3b and Extended Data Fig. 3c)**.

Another key difference is the context of viral infection. Viral replication generates long perfect RNA duplexes, which are more potent activators of PKR than imperfect IR*Alus*.^27,36^ Indeed, DBR1 depletion interfered with poly(I:C)-mediated PKR activation in the non-cancerous BJ and HEK-293T cells, but not in the breast cancer cells **(Extended Data Fig. 3d)**. Although IR*Alus*-containing intron lariats are individually weaker activators of PKR than poly(I:C), their pronounced accumulation upon DBR1 depletion in these cells is most likely sufficient to compensate for the reduced PKR activation by poly(I:C). Furthermore, DBR1 depletion did not downregulate G3BP1, a key stress granule component, in these breast cancer cell lines, further supporting the differential responses to poly(I:C) stimulation **(Extended Data Fig. 3e)**. Based on these, we continued to use MDA-MB-231 to investigate the immunostimulatory effects of intron lariat RNAs.

### PKR preferentially interacts with intron lariat RNAs harboring IR*Alus*

To identify intron lariats that can strongly interact with and activate PKR, we developed an in vitro PKR binding assay followed by RNA-seq (in vitro PKR binding-seq) **(Fig. 3g)**. Briefly, RNase R-treated total RNA from DBR1-depleted or control MDA-MB-231 cells was incubated with recombinant PKR. The RNA-PKR complex was immunoprecipitated and analyzed with input RNA as a control. We found that most PKR-interacting RNAs in siDBR1-transfected cells were derived from introns, whereas intergenic RNAs were most enriched in the control, supporting that DBR1 depletion shifts the PKR-bound RNA pool toward intronic species **(Fig. 3h)**. Additionally, compared to other TEs, the proportion of reads mapped to *Alus* was significantly increased in the PKR-bound RNA **(Extended Data Fig. 3f)**. Moreover, intron RNAs harboring single or multiple SINEs (*Alu* and *MIR)* and LINEs (L1 and L2) were significantly enriched over introns lacking those TEs (classified as ‘Others’), with *Alu*-containing intron RNAs exhibiting the highest PKR binding affinities **(Fig. 3i)**.

Encouraged by these results, we further classified *Alu* associations as direct (DR*Alus*) or inverted (IR*Alus*) based on the relative orientation of neighboring *Alus* within the same intron **(Extended Data Fig. 3g)**. Interestingly, DR*Alus* or single *Alu-*containing RNAs showed high PKR enrichment, while IR*Alus*-containing RNAs demonstrated the strongest PKR binding **(Fig. 3i)**. In addition, having multiple IR*Alus* did not significantly enhance PKR binding compared with a single pair **(Fig. 3i)**. IR*Alus*-PKR binding was not affected by L1 co-existence and intron length, indicating that a single IR*Alus* is sufficient for strong interaction with PKR **(Extended Data Fig. 3h, i)**. We further validated these analyses through RT-qPCR following in vitro PKR binding and pull-down **(Extended Data Fig. 3j)**. Importantly, the enrichment of PKR-intron RNA interaction was abolished when we used the dsRNA-binding domain (dsRBD) mutant (S59A K60A T149A K150A, denoted as PKR dsRBD mutant) **(Extended Data Fig. 3j)**. For the validation experiment, we used FLAG-tagged PKR proteins (both WT and dsRBD mutant) purified from HEK-293T after expressing plasmids encoding these proteins, immunoprecipitating with an anti-FLAG antibody, and eluting. The transcriptome analysis across multiple cell lines showed that in MDA-MB-231, IR*Alus*, single *Alu,* and no TE-containing intron RNAs all showed increased expression upon DBR1 depletion, while in other cell lines, such increase was modest, suggesting that the elevation of IR*Alus*-containing intron RNAs may contribute to robust PKR activation in MDA-MB-231 **(Fig. 3j)**.

To directly assess whether IR*Alus*-containing intron lariats activate PKR, we employed the Complete self-splicing Intron for RNA Circularization (CIRC) method^37^ and synthesized lariat-mimic circular intron RNAs for *DKK1* intron 1 (no TE), *CANX* intron 5, and *PTBP1* intron 13 (both IR*Alus*) **(Fig. 3k and Extended Data Fig. 4a, b; see Methods**). Consistent with our model, both lariat-mimic circular *CANX* intron 5 and *PTBP1* intron 13 significantly activated PKR in vitro, whereas *DKK1* intron 1 did not **(Fig. 3k)**. Furthermore, transfection of circular *CANX* intron 5 into MDA-MB-231 and HEK-293T induced significantly greater PKR activation compared to circular *DKK1* intron 1 **(Fig. 3l)**. In particular, transfection results of *CANX* intron 5 in HEK-293T indicate that increased expression of IR*Alus*-containing intron RNAs can activate PKR.

Next, we investigated the importance of the lariat structure on recognition by PKR through an in vitro debranching assay on RNAs extracted from DBR1-depleted MDA-MB-231 **(Extended Data Fig. 5a)**. We incubated the extracted RNAs with either purified FLAG-tagged DBR1 WT or H85N. Incubation with DBR1 WT resulted in a reduced ratio of lariats to total intron RNAs compared to DBR1 H85N, confirming the successful linearization of lariat RNAs **(Extended Data Fig. 5b)**. We then performed RNA-seq following in vitro PKR-binding using debranched RNAs (DBR1 WT-incubated RNAs). These RNAs exhibited a marked reduction in lariat-derived reads and increased mapping to 5′ splice sites, which are typically excluded during lariat RNA alignment **(Extended Data Fig. 5c, d)**. We found that intron RNAs lacking TEs were less bound by PKR after debranching, suggesting that circularization may generate secondary structures conducive to PKR sensing, as suggested by previous studies **(Extended Data Fig. 5e)**.^38,39^ The overall binding affinity of IR*Alus*-containing RNAs to PKR was also reduced after debranching **(Extended Data Fig. 5e)**. Particularly, as the length of intron RNAs with a single IR*Alus* increases, their interaction with PKR tends to decrease **(Extended Data Fig. 5f)**, suggesting that the lariat structure may promote the pairing between distant IR*Alus*, thereby enhancing PKR recognition. We further found that debranched RNAs exhibited weaker dsRNA signals and reduced PKR activation in vitro **(Extended Data Fig. 5g, h)**. Consistent with these results, lariat-mimic circular *CANX* intron 5 harboring a single IR*Alus* more potently activated PKR in vitro and in cells compared to its linear counterpart **(Fig. 3m and Extended Data Fig. 5i)**.

### PKR-interacting intron lariat RNAs are retained in the nucleus

Given that PKR is primarily localized in the cytoplasm, we examined the nucleo-cytoplasmic distribution of intron lariat RNAs to better characterize their potential interaction with PKR in cells. We fractionated cells and quantified intron RNA levels in the cytoplasmic fraction relative to total RNA using RNA-seq. To our surprise, intron lariat RNAs with high in vitro PKR binding affinity were less localized in the cytoplasm **(Fig. 4a)**. In particular, intron RNAs harboring either IR*Alus* or L1 exhibited high nuclear localization, and the co-presence of both elements further enhanced their nuclear retention **(Fig. 4a, b and Extended Data Fig. 6a)**. Given that longer intron RNAs tend to be more nuclear **(Extended Data Fig. 6b)**, we binned the introns with similar lengths. We found that introns with TEs showed higher nuclear retention across all bins, indicating that both TE presence and intron length jointly regulate RNA subcellular localization **(Fig. 4c)**. We experimentally confirmed the nuclear retention of intron RNAs harboring IR*Alus* and/or L1 through RNA-FISH and RT-qPCR following subcellular fractionation **(Fig. 4d, e)**.

**Fig. 4.**
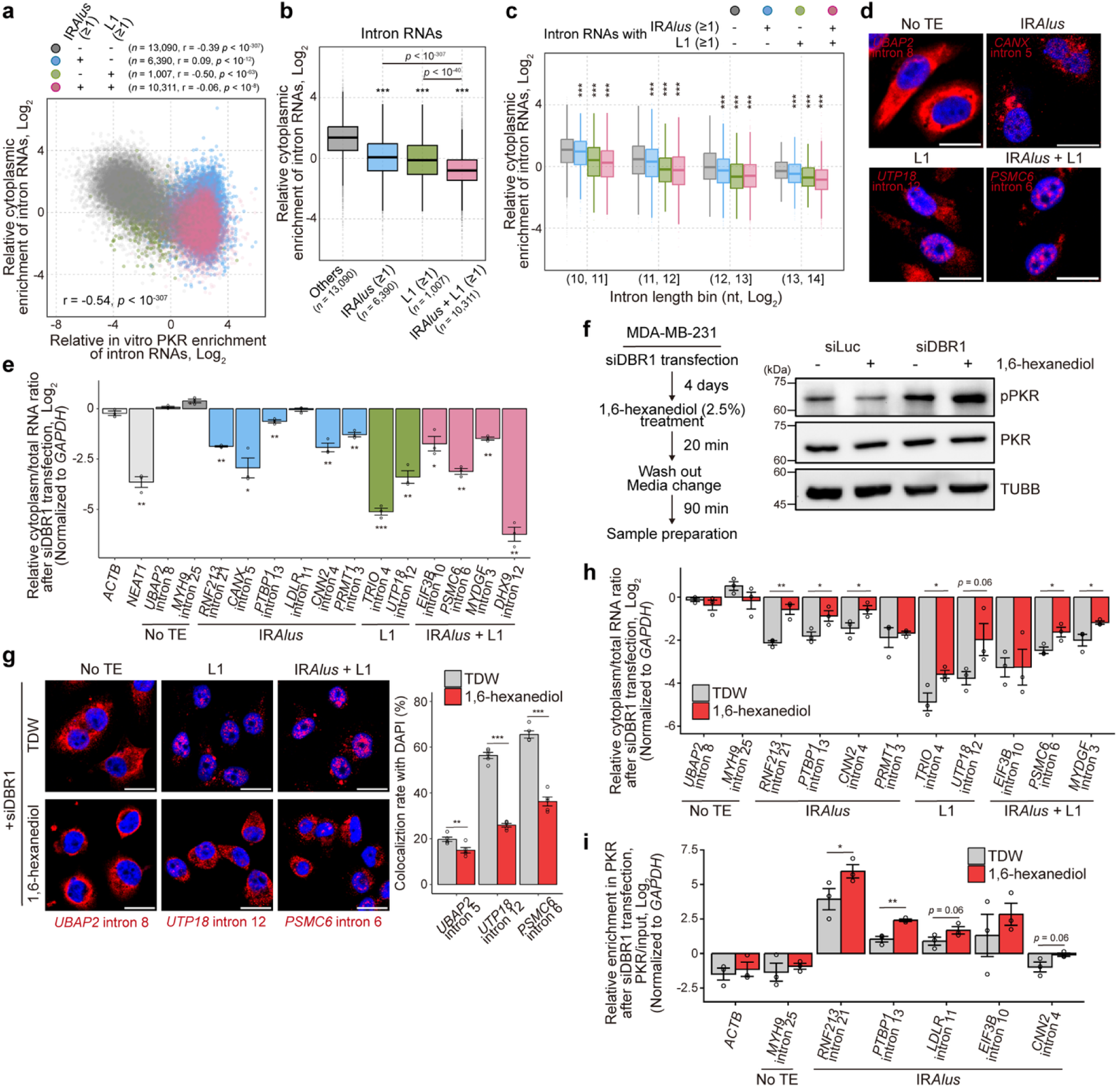
Nuclear retention of TE-containing intron RNAs. **a.** Scatter plot of in vitro PKR enrichment vs. cytoplasmic enrichment (cytoplasm/total RPM ratio) of intron RNAs from MDA-MB-231 6 days after siDBR1 transfection. Each point represents a single intron RNA, categorized based on the presence of either IR*Alus* or L1. **b, c.** Relative cytoplasmic enrichment of intron RNAs with IR*Alus*, L1, or both (**b**) and after binning them based on the length (**c**). RNAs lacking *Alu* and L1 are categorized as “Others”. **d.** RNA-FISH analysis of intron RNAs in DBR1-deficient MDA-MB-231. Blue DAPI signals mark nuclei. Scale bar indicates 20 μm. **e.** Cytoplasm/total ratios of intron RNAs analyzed by RT-qPCR following subcellular fractionation in DBR1-deficient MDA-MB-231 (*n* = 3). *NEAT1* RNA is used as a nuclear control. **f.** WB of PKR activation upon 1,6-hexanediol treatment in MDA-MB-231 transfected with siRNAs for 4 days. **g, h.** Effect of 1,6-hexanediol treatment on the cytoplasmic localization of intron RNAs analyzed by RNA-FISH (**g**, *n* = 5) and RT-qPCR following subcellular fractionation (**h**, *n* = 3) in MDA-MB-231 transfected with siDBR1 for 4 days. Co-localization rate between red and blue signals is shown on the right (**g**). Scale bar indicates 20 μm. **i.** RT-qPCR of selected intron RNAs pulled down with PKR after 1,6-hexanediol treatment in DBR1-deficient MDA-MB-231, normalized to the input (*n* = 3). All error bars denote s.e.m. *P*-values were calculated using either the two-sided Wilcoxon signed-rank test (b, c) or the one-sided Student’s t-test (e, g, h, i); ^∗^*p* < 0.05, ^∗∗^*p* < 0.01, ^∗∗∗^*p* < 0.001. Correlation *r* and its *p*-value were calculated using the Spearman correlation (a).

The nuclear retention of RNAs with high PKR binding potential raises questions about their in-cellulo interactions with PKR. First, we treated cells with 1,6-hexanediol, a small molecule commonly used to reversibly disrupt the nuclear pore complex and elicit nuclear content leakage, to examine the effect of releasing nucleus-retained RNAs.^40^ Notably, 1,6-hexanediol treatment increased pPKR in DBR1 knockdown cells but not in control cells **(Fig. 4f)**. Indeed, many nucleus-retained intron lariat RNAs were released into the cytoplasm upon 1,6-hexanediol treatment **(Fig. 4g, h)**. Correspondingly, their cytoplasmic release led to increased interaction with PKR, confirmed by RT-qPCR following PKR fCLIP, suggesting that nucleus-retained intron RNAs can serve as PKR activators **(Fig. 4i)**.

### Enhanced PKR-intron lariat RNA interaction during mitosis elicits PKR hyperactivation

Since the nuclear membrane naturally disintegrates during mitosis, we investigated the role of the cell cycle in mediating cellular response to DBR1 depletion. Indeed, nuclear intron lariat RNAs are released upon M-phase arrest, increasing the cytoplasmic dsRNA signal while decreasing the nuclear signal **(Fig. 5a and Extended Data Fig. 6c)**. In contrast, when cells were arrested at S-phase, cytoplasmic levels of TE-containing intron RNAs were significantly reduced **(Fig. 5b)**.

**Fig. 5.**
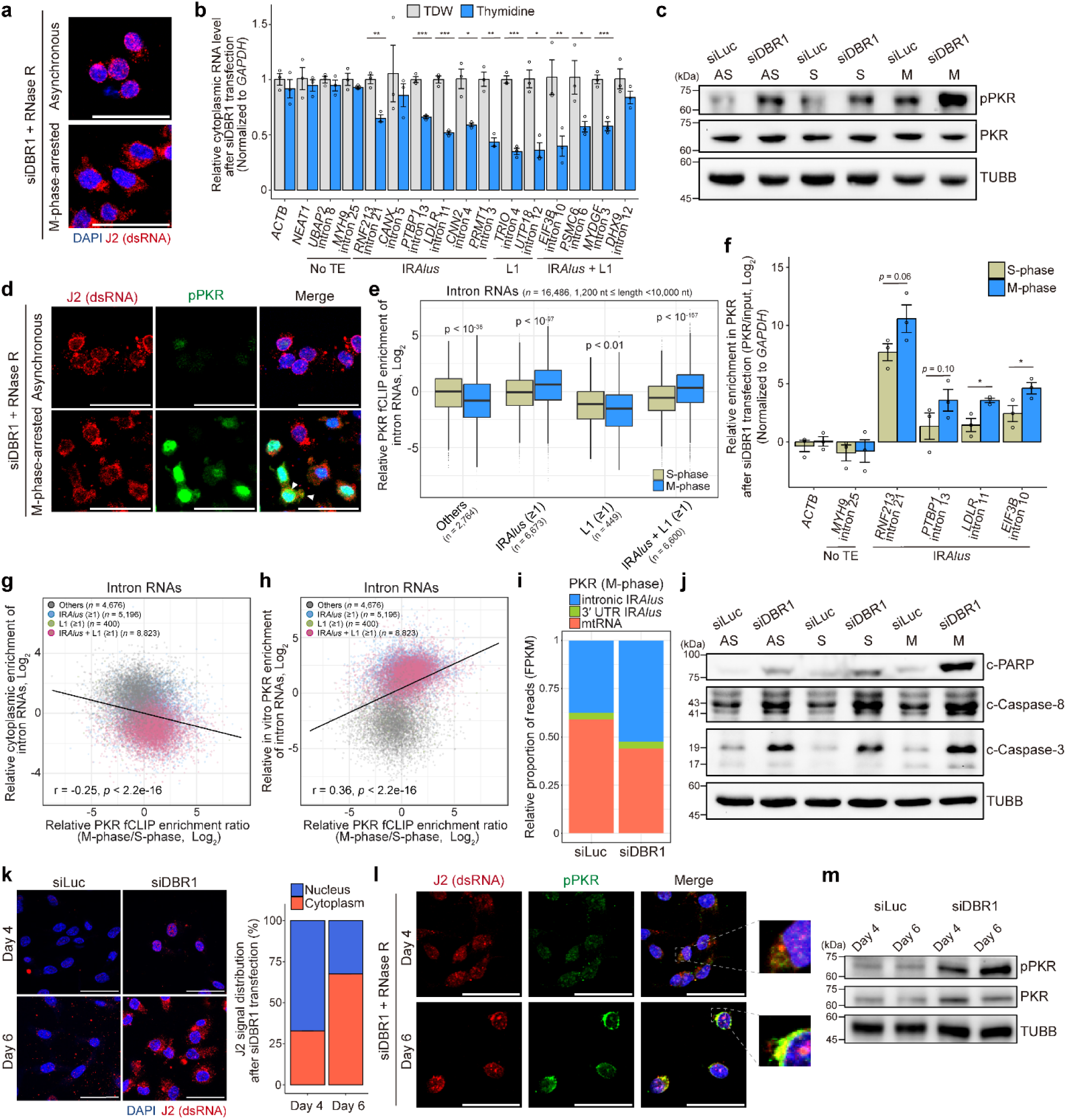
Enhanced intron lariat RNA-PKR interactions during mitosis. **a.** J2 signals following RNase R treatment on M-phase-arrested DBR1-deficient MDA-MB-231. **b.** Relative cytoplasmic levels of intron RNAs upon thymidine treatment analyzed by RT-qPCR following subcellular fractionation in DBR1-deficient MDA-MB-231 (*n* = 3). **c.** WB of PKR phosphorylation after S-or M-phase arrest in DBR1-deficient MDA-MB-231; AS: asynchronous, S: S-phase-arrested, M: M-phase-arrested. **d.** J2 and pPKR signals following RNase R treatment on M-phase-arrested DBR1-deficient MDA- MB-231. White arrows indicate the co-localization of PKR and dsRNAs. **e.** Relative PKR enrichment of intron RNAs in PKR fCLIP-seq (PKR/input RPM ratio) on S-or M-phase-arrested DBR1-deficient MDA-MB-231, categorized by the presence of IR*Alus*, L1, or both. RNAs lacking *Alu* and L1 are categorized as “Others”. **f.** RT-qPCR of selected intron RNAs pulled down with PKR on S-or M-phase-arrested DBR1- deficient MDA-MB-231, normalized to the input (*n* = 3). **g, h.** Scatter plot of PKR fCLIP enrichment ratio (M-phase/S-phase PKR enrichment ratio) vs. cytoplasmic enrichment (cytoplasm/total RPM ratio) (**g**) or in vitro PKR enrichment (**h**) of intron RNAs from MDA-MB-231. Each point represents a single intron RNA, categorized based on the presence of either IR*Alus* or L1. **i.** Relative proportion of FPKM values corresponding to intronic IR*Alus*, 3′ UTR IR*Alus*, and mtRNAs bound to PKR on M-phase-arrested siRNA-transfected cells. **j.** WB of apoptotic markers on S-or M-phase-arrested DBR1-deficient MDA-MB-231. **k.** J2 signals 4 and 6 days after siRNA transfection to MDA-MB-231. The quantified average percentage of nuclear and cytoplasmic J2 signals is shown at the right (*n =* 10). **l, m.** Immunocytochemistry for J2 and pPKR signals following RNase R treatment (**l**) and WB for pPKR (**m**) 4 and 6 days after siDBR1 transfection to MDA-MB-231. All error bars denote s.e.m. All scale bars indicate 50 μm. *P*-values were calculated using either the one-sided Student’s t-test (b, f) or the two-sided Wilcoxon signed-rank test (e); ^∗^*p* < 0.05, ^∗∗^*p* < 0.01, ^∗∗∗^*p* < 0.001. Correlation *r* and its *p*-value were calculated using the Spearman correlation (g, h).

If nuclear TE-containing intron lariat RNAs are released to the cytoplasm during mitosis, PKR activation by DBR1 depletion should exhibit a cell cycle-dependent pattern. Indeed, PKR was hyperactivated in M-phase-arrested DBR1-deficient cells compared to that of S-phase-arrested and asynchronous (AS) cells, accompanied by increased co-localization with intronic dsRNAs **(Fig. 5c, d and Extended Data Fig. 6d)**. We then performed PKR fCLIP-seq on M- or S-phase-arrested DBR1-deficient cells and found increased interaction between IR*Alus*-containing intron RNAs and PKR in M-phase, which was validated by RT-qPCR **(Fig. 5e, f)**. Interestingly, RNAs showing strong enhancement in PKR interaction in M-phase were more retained in the nucleus in interphase and showed high PKR binding in vitro, indicating that nuclear retention may limit PKR activation in DBR1-deficient cells **(Fig. 5g, h)**. In terms of the relative composition of major PKR substrates during mitosis, we found that intron RNAs became the dominant PKR substrate in DBR1-deficient cells, whereas mtRNAs were the dominant species in control cells **(Fig. 5i)**. PKR hyperactivation was accompanied by enhanced induction of apoptotic markers and increased peIF2α **(Fig. 5j and Extended Data Fig. 6e)**, which was abolished in PKR KO cells, proposing that PKR hyperactivation accelerates mitotic cell death in DBR1-deficient cells **(Extended Data Fig. 6f).**

Consistent with our model, the dsRNA signal probed by the J2 antibody was initially high in the nucleus 4 days after DBR1 knockdown, but shifted to the cytoplasm 6 days post-knockdown **(Fig. 5k)**. This was accompanied by PKR co-localization with dsRNAs in the cytoplasm and enhanced PKR activation **(Fig. 5l, m)**. Moreover, PKR hyperactivation persisted longer in DBR1-deficient cells than in control after release from mitotic arrest **(Extended Data Fig. 6g-i)**. Notably, cytoplasmic intron lariat RNAs harboring IR*Alus* showed higher stability than that of 3′ UTR IR*Alus* **(Extended Data Fig. 6j)**. This superior stability of intron lariats may account for prolonged PKR activation upon release from mitotic arrest in DBR1-deficient cells.

### DBR1 depletion enhances the sensitivity to anti-mitotic drugs in vitro and in vivo

Considering the highly proliferative nature of cancer cells and the mechanism of anti-mitotic chemotherapy drugs that kill cancer cells via M-phase arrest,^41,42^ we investigated the potential synergistic effect of combining DBR1 inhibition with anti-mitotic drugs for cancer therapy, such as paclitaxel (PTX) and ixabepilone. Aligned with our model, DBR1 depletion sensitized MDA-MB-231 to PTX **(Fig. 6a)**. Moreover, this combination resulted in significantly higher pPKR and c-PARP than depletion/treatment of either one alone **(Fig. 6b, c and Extended Data Fig. 7a)**. Importantly, the enhanced sensitivity to PTX was nearly abolished in PKR KO cells **(Fig. 6d, e)**.

**Fig. 6.**
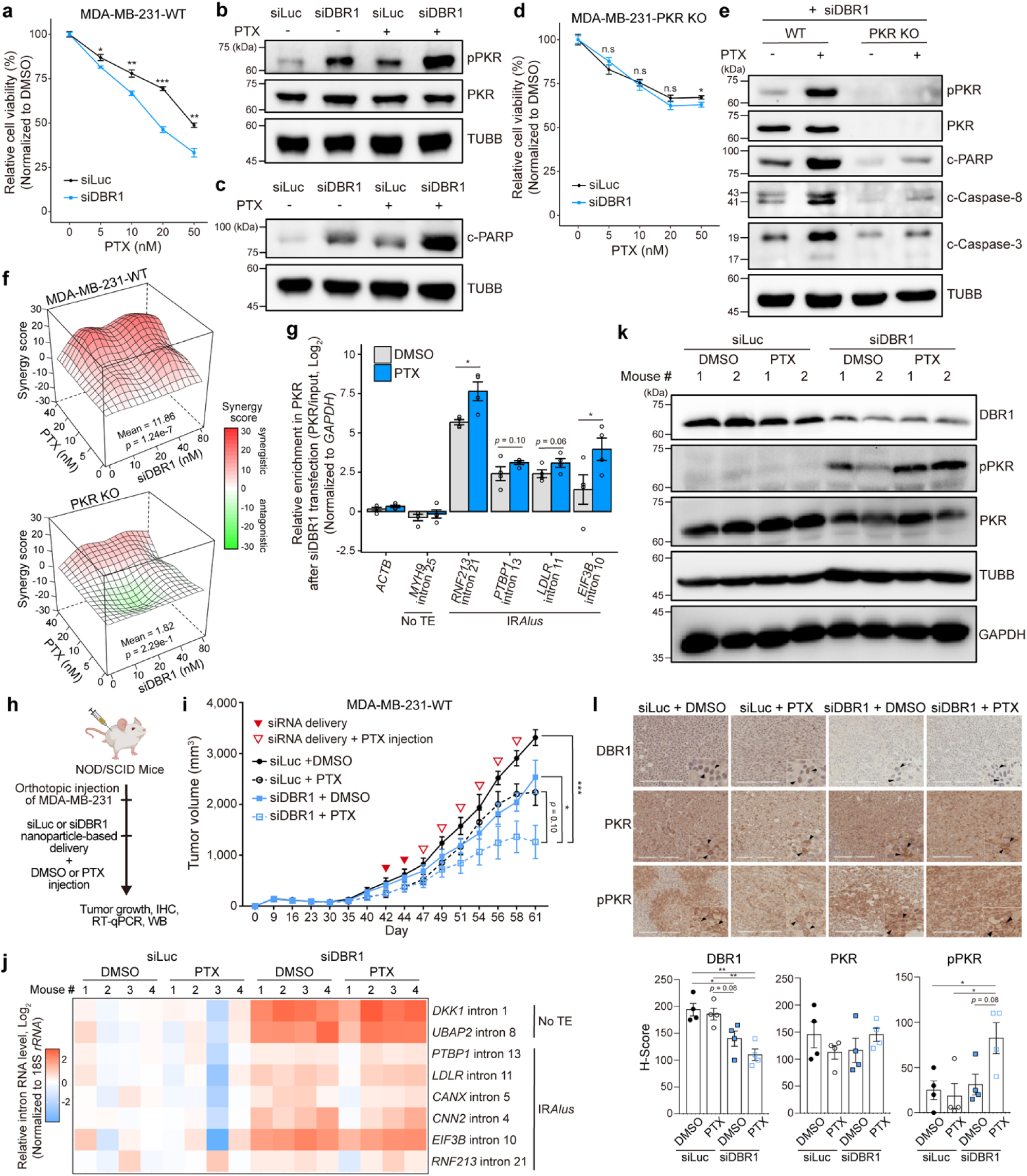
Enhanced sensitivity to anti-mitotic drugs upon DBR1 depletion. **a.** Cell viability upon the treatment of PTX (*n* = 4 for each concentration) in siRNA-transfected MDA-MB-231. For both siRNAs, cell viabilities were normalized to their respective no-PTX treatment condition. **b, c.** WB for PKR phosphorylation (**b**) and PARP cleavage (**c**) upon 20 nM PTX treatment in siRNA-transfected MDA-MB-231. **d.** Cell viability upon the treatment of PTX (*n* = 4 for each concentration) in siRNA-transfected MDA-MB-231 PKR KO. **e.** WB of apoptotic markers upon 20 nM PTX treatment in siDBR1-transfected MDA-MB-231 WT and PKR KO. **f.** Synergy scores between siDBR1 and PTX in MDA-MB-231 WT and PKR KO, calculated using the Bliss independence model. Mean synergy scores and p-values are indicated. Positive and negative scores indicate synergistic and antagonistic interactions, respectively. **g.** RT-qPCR analysis of selected intron RNAs pulled down with PKR upon 20 nM PTX treatment in siDBR1-transfected MDA-MB-231, normalized to the input (*n* = 4). **h.** Experimental scheme of an in vivo mouse experiment. Nanoparticle–mediated delivery of siRNA was followed by DMSO or PTX treatment in tumor sites derived from orthotopic injection of MDA-MB-231 into NOD/SCID mice. Tumor volume was monitored at defined time points, and tumor tissues were collected after sacrifice for immunohistochemical staining, RT-qPCR, and WB. **i.** Tumor growth curves of orthotopic MDA-MB-231 WT xenografts in NOD/SCID mice treated with nanoparticle-formulated siRNA, followed by DMSO or PTX intraperitoneal administration. Red-filled arrowheads indicate the timing of siRNA delivery, and open arrowheads denote siRNA delivery combined with PTX injection. Tumor volumes were measured at the indicated time points and are presented as mean ± s.e.m. (*n* = 4 per group). **j.** Heatmap showing relative intron RNA levels in tumor tissues from NOD/SCID mice treated with nanoparticle-formulated siLuc or siDBR1, followed by DMSO or PTX administration, quantified by RT-qPCR (*n* = 4 per group). **k, l.** WB (**k**) and immunohistochemistry (**l**) of DBR1 expression and PKR phosphorylation in tumor tissues from NOD/SCID mice treated with nanoparticle-formulated siRNA, followed by DMSO or PTX administration. All error bars denote s.e.m. *P*-values were calculated using the one-sided Student’s t-test (a, d, g) and Tukey’s HSD test following a one-way ANOVA (i, l); ^∗^*p* < 0.05, ^∗∗^*p* < 0.01, ^∗∗∗^*p* < 0.001, and n.s: non-significant.

To further assess the combinatorial effects, we performed a drug synergy analysis utilizing the Bliss independence model across varying concentrations of PTX and siDBR1. We first confirmed the increase in DBR1 knockdown as siDBR1 concentration increased **(Extended Data Fig. 7b)**. The combination yielded a strongly synergistic response in WT cells, which was significantly reduced in PKR KO cells and siLuc-transfected cells **(Fig. 6f and Extended Data Fig. 7c)**. DBR1 depletion also considerably increased the sensitivity to ixabepilone in a PKR-dependent manner, ruling out PTX-specific effects **(Extended Data Fig. 7d-g)**. Furthermore, we confirmed increased sensitivity to PTX upon DBR1 depletion in MCF7 **(Extended Data Fig. 7h, i)**. Of note, our PKR fCLIP experiment showed that PTX treatment increased the binding of intron RNAs harboring IR*Alus* to PKR, accounting for the increased efficacy of these drugs **(Fig. 6g)**.

Next, we evaluated the combined effect of DBR1 depletion and anti-mitotic drugs in vivo. We first orthotopically injected MDA-MB-231 cells into NOD/SCID mice. Once tumors reached a measurable size, we administered intratumoral injections of siDBR1 or siLuc complexed with a nanoparticle-based in vivo delivery reagent, followed by PTX treatment, and subsequently monitored tumor volume **(Fig. 6h)**. First, we confirmed that siDBR1 injection alone had no effect on mouse body weight, whereas PTX treatment tended to slightly decrease it, thereby ruling out potential adverse effects of siDBR1 **(Extended Data Fig. 7j)**. The combination of siDBR1 delivery and PTX treatment suppressed tumor growth more than either treatment alone, although statistical support was modest owing to inter-individual variation and the small sample size **(Fig. 6i)**. We then extracted RNA and protein from tumor tissues and confirmed DBR1 downregulation, accompanied by concomitant accumulation of intron RNA, in siDBR1-injected tumors **(Fig. 6j, k)**. In addition, western blotting and immunohistochemistry revealed robust PKR activation in tumors treated with both siDBR1 and PTX **(Fig. 6k, l)**. Together, these results suggest that DBR1 depletion sensitizes breast cancer cells to anti-mitotic drugs through PKR hyperactivation in vitro, with a consistent trend in vivo.

### RNA editing in intronic IR*Alus* mitigates PKR activation

Cells employ adenosine-to-inosine (A-to-I) editing, catalyzed by adenosine deaminase acting on RNA 1 (ADAR1), to prevent immune activation by self-dsRNAs.^43–45^ Recent comprehensive analyses have revealed that *Alu*s are primary editing substrates,^45,46^ prompting us to examine whether accumulated intron lariat RNAs undergo A-to-I editing. We identified editing sites and measured editing rates by detecting A-to-G mismatches in RNA-seq reads aligned to intronic regions from siDBR1-transfected MDA-MB-231 **(Fig. 7a)**. Our analysis revealed that IR*Alus*-containing intron RNAs exhibited high A-to-I editing scores (as defined in **Fig. 7a**) compared to those without IR*Alus* **(Fig. 7b)**. At the same time, intron RNAs showed low C-to-U editing scores regardless of the presence of IR*Alus* **(Extended Data Fig. 8a)**. Interestingly, intron RNAs harboring either *Alu* or DR*Alus* showed slightly higher A-to-I editing scores than those lacking *Alu* but lower than those with IR*Alus* **(Extended Data Fig. 8b)**. Furthermore, more than 90% of A-to-I editing events in intron RNAs containing IR*Alus* occur within *Alu* repeats (either sense or antisense *Alu*) **(Fig. 7c)**. Lastly, A-to-I edited IR*Alus*-containing RNAs harbor a greater number of IR*Alus* than unedited counterparts, suggesting that an increased likelihood of *Alu* duplex formation resulting from the higher *Alu* content may promote editing **(Extended Data Fig. 8c)**.

**Fig. 7.**
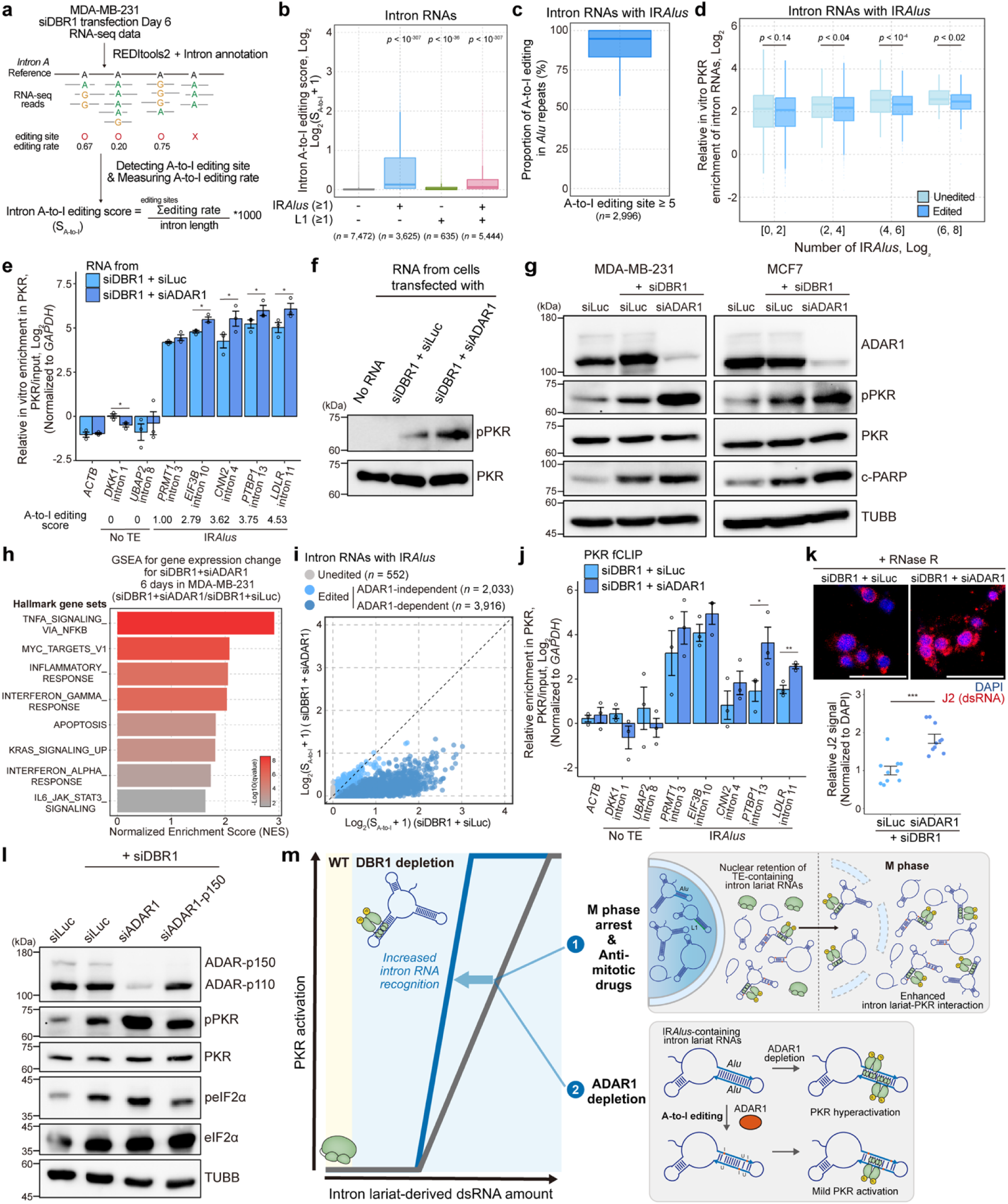
A-to-I editing in IR*Alus*-containing intron lariat RNAs. **a.** Schematic of the A-to-I editing site identification and editing score calculation. **b.** A-to-I editing scores of intron lariat RNAs depending on the presence of either IR*Alus* or L1. **c.** Proportion of A-to-I editing on *Alu*s (either sense or antisense *Alu*) of intron RNAs containing IR*Alus.* RNAs with at least five editing sites are included in the analysis. **d.** Comparison of relative in vitro PKR enrichment for A-to-I edited and unedited IR*Alus*-containing intron RNAs, categorized by the number of IR*Alus*. **e.** RT-qPCR of intron RNAs following in vitro PKR binding assay using total RNAs extracted from siDBR1-transfected or siDBR1/siADAR1-co-transfected MDA-MB-231, normalized to input (*n* = 3). **f.** In vitro PKR activation assay using recombinant PKR and total RNAs extracted from MDA-MB- 231 transfected with siDBR1 alone or co-transfected with siDBR1 and siADAR1. The mass ratio of RNA to PKR is 1:1. **g.** Effect of ADAR1 depletion on PKR phosphorylation and PARP cleavage in DBR1-deficient MDA-MB-231 and MCF7. **h.** Hallmark gene sets significantly enriched in MDA-MB-231 co-transfected with siDBR1 and siADAR1 compared to siDBR1-only, as identified by GSEA. **i.** Changes in A-to-I editing scores of IR*Alus*-containing intron RNAs after ADAR1 depletion in DBR1-deficient MDA-MB-231. Each point represents a single intron RNA. Intron RNAs showing ≥ 50% reduction in A-to-I editing score upon ADAR1 depletion are classified as “ADAR1-dependent”. **j.** RT-qPCR of selected intron RNAs pulled down with PKR upon ADAR1 depletion in siDBR1-transfected MDA-MB-231, normalized to the input (*n* = 3). **k.** Effect of ADAR1 depletion on J2 signal following RNase R treatment in DBR1-deficient MDA- MB-231. Quantification is shown below (*n* = 10). Scale bar indicates 50 μm. **l.** Effect of the ADAR1-p150 depletion on PKR and eIF2α phosphorylation in DBR1-deficient MDA-MB-231. **m.** A model for the immunoregulation by intron lariat RNAs. In DBR1-deficient cells, intron lariat-derived dsRNAs accumulate, but are largely retained in the nucleus and undergo A-to-I editing, which prevents their recognition by PKR in cells. Induction of mitotic arrest by anti-mitotic drugs or depletion of ADAR1 enhances intron lariat-PKR interaction, leading to PKR hyperactivation and increased cell death. All error bars denote s.e.m. *P*-values were calculated using either the two-sided Wilcoxon signed- rank test (b, d) or the one-sided Student’s t-test (e, j, k); ^∗^*p* < 0.05, ^∗∗^*p* < 0.01, ^∗∗∗^*p* < 0.001.

We then examined how A-to-I editing affects PKR binding. We binned intron RNAs by the number of IR*Alus* and found that A-to-I edited intron RNAs containing IR*Alus* bound PKR more weakly than unedited ones **(Fig. 7d)**. We then performed an in vitro PKR binding assay using total RNAs from siDBR1-and siADAR1-co-transfected cells and found that among IR*Alus*-containing RNAs, those with high A-to-I editing scores exhibited enhanced binding to PKR upon ADAR1 depletion **(Fig. 7e)**. Furthermore, RNAs from siDBR1 and siADAR1-co-transfected cells activated PKR in vitro more effectively than those from siDBR1-transfected cells alone **(Fig. 7f)**. Consistent with these results, dual depletion of ADAR1 and DBR1 triggered a significant increase in PKR activation and cell death in both MDA-MB-231 and MCF7 **(Fig. 7g and Extended Data Fig. 8d)**. In addition, our RNA-seq analysis revealed that additional ADAR1 depletion significantly upregulates innate immune-related pathways, including TNFα signaling and IFN responses, in DBR1-deficient cells **(Fig. 7h)**. Consistently, we found that most IR*Alus*-containing intron RNAs, such as *PTBP1* intron 13 and *LDLR* intron 11, exhibited reduced A-to-I editing upon ADAR1 depletion **(Fig. 7i and Extended Data Fig. 8e)**. Moreover, highly edited intron RNAs show increased PKR enrichment upon ADAR1 depletion in DBR1-deficient cells **(Fig. 7j)**. Lastly, ADAR1 depletion substantially elevated the levels of intron lariat-derived dsRNAs, indicating that ADAR1-mediated A-to-I editing may reduce the duplex integrity of intron lariats **(Fig. 7k)**.

Given that ADAR1 is expressed as two isoforms -a constitutively expressed short nuclear p110 and a long IFN-inducible cytoplasmic p150, we investigated their relative contribution and found that ADAR1-p150 depletion was insufficient to trigger PKR activation and cell death **(Fig. 7l and Extended Data Fig. 8f)**. ADAR1-p150 depletion partially reduced, or in some cases did not reduce, A-to-I editing **(Extended Data Fig. 8e)**. Our results align with recent studies where ADAR1-p150-selective editing sites are enriched in 3′ UTR, and upon ADAR1-p150 loss, residual editing sites are predominantly found in introns.^47,48^ Given the low basal ADAR1-p150 expression relative to ADAR1-p110 in DBR1-deficient cells **(Fig. 7l)** and the nuclear localization of IR*Alus*-containing intron RNAs, A-to-I editing of intron lariats is likely mediated primarily by ADAR1-p110.

Lastly, we analyzed the impact of A-to-I editing on the subcellular localization of intron RNAs. Interestingly, when binned by the number of IR*Alus*, A-to-I edited RNAs were more localized in the cytoplasm compared to unedited ones, explaining why ADAR1 depletion can significantly enhance PKR sensing of highly edited intron RNAs **(Extended Data Fig. 8g)**. To assess whether A-to-I editing directly influences their localization, we performed cytoplasmic and total RNA-seq and compared the relative cytoplasmic enrichment of intron RNAs following ADAR1 depletion in DBR1-deficient cells. Unexpectedly, intron RNAs with a markedly reduced editing upon ADAR1 depletion (ADAR1-dependent) exhibited no substantial change in their cytoplasmic enrichment compared to ADAR1-independent or unedited intron RNAs, regardless of the number of IR*Alus* **(Fig. 7i and Extended Data Fig. 8h-j)**. These results suggest that A-to-I editing has a minimal direct impact on the subcellular localization of intron RNAs, indicating that other factors likely drive their cytoplasmic enrichment.

## DISCUSSION

Our study uncovers undegraded intron lariat RNAs as self-immunogens, revealing that TE- containing RNAs, particularly those with IR*Alus*, can directly activate PKR, and cells counteract this through nuclear sequestration and RNA editing. Importantly, we can exploit this mechanism by combining DBR1 depletion with anti-mitotic drugs or ADAR1 knockdown as effective cancer therapeutics **(Fig. 7m)**.

Over the past decades, multiple studies have demonstrated that the depletion or mutation of dsRNA regulators, such as *DICER1* and *ADAR*, leads to aberrant innate immune activation and contributes to the pathogenesis of inflammatory and autoimmune diseases.^49–51^ However, recent studies have shown that DBR1 deficiency can impair exogenous dsRNA-mediated antiviral responses by disrupting stress granule assembly and promoting the cytoplasmic accumulation of intronic IR*Alus*-derived dsRNAs that titrate PKR.^24–27^ Our data indicate that the extent of intronic IR*Alus*-derived dsRNA accumulation may be a key determinant of whether DBR1 depletion activates PKR. Cells with a high IR*Alus* burden (breast cancer), sustain PKR activation that overrides titration-mediated suppression, whereas cells with low accumulation (HEK-293T and BJ), do not. The preserved G3BP1 levels in the analyzed breast cancer cells may further allow them to evade DBR1 deficiency-induced PKR inhibition. Unraveling the molecular mechanism underlying the preferential upregulation of TE-containing introns will enhance our understanding of the effects of DBR1 deficiency.

Our study reveals the physical and chemical mechanisms that prevent non-self-sensing of intron lariat RNAs. Physically, intron lariat RNAs harboring either IR*Alus* or L1 exhibited reduced cytoplasmic localization. Previous studies have shown that mRNAs containing IR*Alus* in 3′ UTR are sequestered in paraspeckles.^52,53^ This can be reversed by STAU1, but it also protects IR*Alus* from PKR recognition.^54^ Our study adds to the current literature by showing that TE-harboring intron lariat RNAs are also subject to nuclear retention and that co-presence of IR*Alus* and L1 enhances the degree of sequestration. Chemically, A-to-I editing may function as an RNA surveillance mechanism by reducing interaction with PKR. Moreover, given recent findings that ADAR1 can directly inhibit PKR,^55,56^ ADAR1 may use multiple mechanisms to prevent intron lariat RNA-mediated PKR activation beyond its RNA-editing activity. Overall, our findings suggest that multiple cellular mechanisms work in parallel to mitigate the sensing of immunogenic intron lariat RNAs.

Since certain circular RNAs can form short imperfect RNA duplexes that inhibit PKR,^38,39^ some intron lariats may adopt similar secondary structures and exert PKR inhibitory effects. Our PKR fCLIP-seq analysis revealed that PKR interacts with a subset of TE-lacking intron lariats that are predominantly cytoplasmic, suggesting a tug-of-war between PKR-activating IR*Alus*-containing intron lariats and PKR-inhibitory TE-lacking intron lariats, modulated by RNA localization and relative abundance. Notably, this balance shifts spatiotemporally with the cell cycle. During mitosis, nucleus-retained IR*Alus*-containing intron lariats win over cytoplasmic TE-lacking intron lariats in PKR binding. Moreover, upon mitotic exit, IR*Alus*-containing intron lariats are markedly more stable than mRNAs harboring 3′ UTR IR*Alus* in the cytoplasm, implying that DBR1-deficient cells may fail to restore cytosolic dsRNA homeostasis after mitosis.

One notable finding is that intron lariat-derived dsRNAs selectively activate PKR rather than MDA5 and RIG-I, consistent with a prior study showing that intronic inverted retroelement-derived dsRNAs fail to activate RIG-I and MDA5 while effectively engaging PKR.^16^ Structurally, since RIG-I requires a 5′-triphosphate moiety and MDA5 preferentially recognizes long uninterrupted duplexes,^13^ intron lariat-derived edited dsRNAs may evade sensing. In contrast, PKR can recognize even shorter, imperfect duplexes. Additionally, DBR1 depletion in MDA-MB- 231 significantly upregulates PKR expression without equivalently inducing MDA5 and RIG-I, which may favor selective activation of PKR signaling.

Endogenous dsRNAs are emerging targets for cancer immunotherapy due to their ability to induce viral mimicry and enhance anti-tumor responses.^18,20,23^ Our study adds DBR1 depletion as a trigger for intronic dsRNA accumulation. Moreover, we further found that anti-mitotic drugs can facilitate the cytosolic release of intron dsRNAs, thereby amplifying PKR-mediated cancer cell death. Given that TE transcripts preferentially localize in the nucleus,^57^ combinations of anti-mitotic drugs with other TE-reactivating agents may further enhance cytosolic dsRNA accumulation and anti-tumor immunity. In this context, ADAR1 has been identified as a promising therapeutic target in various cancers, particularly breast cancer,^58,59^ and indeed, dual depletion of DBR1 and ADAR1 triggers PKR hyperactivation and increased cancer cell death, presenting a complementary immunotherapeutic strategy. In summary, our study uncovers multi-layered mechanisms that shield cells from innate immune activation by self-intron lariat RNAs and proposes targeting the debranching process as a potential therapeutic strategy to enhance the efficacy of anti-mitotic drugs.

## ACKNOWLEDGEMENTS

We appreciate Dr. Chanhee Kang for providing BJ cells. This research was supported by the Bio&Medical Technology Development Program of the National Research Foundation (NRF) funded by the Korean government’s Ministry of Science and ICT (RS-2024-00440866 and RS-2025-16066141) and funded by the Korean government’s Ministry of Health & Welfare and Ministry of Science and ICT (RS-2025-25458777). K.L. was supported by the KAIST Jang Young Sil Fellow Program.

## AUTHOR CONTRIBUTIONS

K.L., J.K., H.S.R., and Y.K. designed the study and analysis. In vitro and cell experiments were performed by K.L., J.K., T.A.W., M.J., N.K., H.C., and M.J.K.. In vivo experiments were performed by S.Y.P., D.S.K., H.Y.K., H.C.. Bioinformatic analysis was performed by K.L., J.K., and W.D.J.. The study was supervised by H.S.R. and Y.K.. K.L. and Y.K. wrote the manuscript with the contributions from J.K., S.Y.P., and H.S.R.. All authors reviewed the data and edited the manuscript.

## SUPPLEMENTAL INFORMATION

Document S1. Extended Data Figs 1-8 and Tables 1-5

## METHODS

### Cell lines and culture

MDA-MB-231, MCF7, and Hs578T cells were grown in Roswell Park Memorial Institute (RPMI) 1640 medium (Welgene, LM011-01) supplemented with 9.1% FBS (Sigma-Aldrich, F2442) and 1% Penicillin-Streptomycin (PS, Welgene, LS202-02). HEK-293T cells were grown in Dulbecco’s Modified Eagle Medium (DMEM, Welgene, LM001-05) supplemented with 9.1% FBS. BJ and IMR-90 cells expressing human telomerase reverse transcriptase (hTERT) were maintained in DMEM supplemented with 15% FBS, 1% PS, and 0.1 mM non-essential amino acids (Welgene, LS005-01). All cell lines were maintained in a 37°C and 5% CO_2_ humidified incubator and passaged at a 1:5 dilution every 3-4 days.

### In vivo mouse experiments

Five-week-old female NOD/SCID mice were obtained from KOATECH (Seoul, Korea). All procedures were approved by the Institutional Animal Care and Use Committee (IACUC) of Seoul National University (Approval No. SNU-231218-3-1) and conducted in accordance with institutional and national animal care guidelines.

To investigate the effects of DBR1 knockdown and paclitaxel (PTX, LC Laboratories, P-9600) on breast cancer progression, an orthotopic xenograft experiment was performed using MDA-MB-231 cells. Briefly, 8 × 10⁶ cells were suspended in a 1:1 mixture of Matrigel (Corning, 354234) and cold PBS and orthotopically injected into the mammary fat pads of mice. Tumor growth was monitored three times weekly using calipers, and mouse body weight was recorded once per week. Tumor volume was calculated using the formula: tumor volume (mm³) = (a × b²)/2, where a represents the longest diameter and b represents the perpendicular diameter.

When tumors reached 100–150 mm³, mice were randomized by tumor burden into four treatment groups (n = 4 per group; 16 mice in total). For each injection, 10 μg of siLuc or siDBR1 was complexed with a nanoparticle-based in vivo transfection reagent (Altogen, 5032) according to the manufacturer’s protocol and administered intratumorally under isoflurane anesthesia three times per week, beginning on day 42 after tumor cell implantation. From day 49, PTX (15 mg/kg) was given intraperitoneally without anesthesia, concomitantly with siRNA treatment. At the study endpoint, mice were euthanized by carbon dioxide inhalation, and tumors were excised, weighed, and fixed in 10% neutral-buffered formalin for histological and immunohistochemical analyses.

### RNA transfection

siRNAs and poly(I:C) (Sigma-Aldrich, P1530) were transfected into cells using Lipofectamine 3000 (Thermo Fisher Scientific, L3000075) following the manufacturer’s protocol. For knockdown of DBR1 for 4 and 6 days, siRNAs were transfected into cells every 2 and 3 days, respectively. As a negative control, siLuc (targeting Luciferase) was used. siRNAs were purchased from Bioneer Korea. The sequences of the siRNAs used in this study are summarized in **Table 1**. For the transfection of RNAs extracted from cells, we used the *Trans*IT-mRNA Transfection Kit (Mirus, MIR2250) following the manufacturer’s protocol.

### shRNA plasmid production and lentivirus transduction

The pLKO.1 plasmid (Addgene, 8453) was digested with AgeI and EcoRI restriction enzymes, followed by purification via gel extraction. shRNA-sequence DNA oligos were annealed in a DNA annealing buffer (10 mM Tris-HCl pH 8.0 and 80 mM NaCl) by heating to 95°C for 5 min and ligated into the cleaved plasmid using T4 DNA ligase at room temperature (RT) for 1 h. The plasmid was transformed into a DH5α competent *E. coli*. Lentivirus particles were produced in HEK-293T cells by co-transfecting pLKO.1-shRNA with psPAX2 (Addgene, 12260) and pMD2.G (Addgene, 12259) supplemented by 1% (v/v) Bovine Serum Albumin (BSA). Media containing lentivirus particles were harvested after 60 h of transfection and transduced into target cells. After 2 days of transduction, puromycin (InvivoGen, ant-pr-1) was added to the media for selection. pLKO.1 scrambled shRNA (shScramble) was used as a negative control. Sequences of the shRNA used in this study are provided in **Table 2**.

### Chemical treatment and cell viability test

To arrest cells at the S-or M-phase, the double-thymidine block method was employed. Briefly, 72 h after siDBR1 transfection, cells were treated with 2 mM thymidine (Sigma-Aldrich, T9250) for 18 h and then released into a fresh medium. After 9 h, 2 mM thymidine (for S-phase arrest) or 100 ng/mL nocodazole (Sigma-Aldrich, M1404, for M-phase arrest) was added to the medium, and cells were incubated for an additional 17 h. For anti-mitotic drug treatment, paclitaxel or ixabepilone (Sigma-Aldrich, SML2615) was added to the medium 72 h post-siRNA transfection, and cells were incubated for an additional 48 h. Synergy scores between siRNA and PTX were calculated using the Bliss independence model. To inhibit apoptosis, 30 μ M Z-VAD-FMK (InvivoGen, tlrl-vad) was added to the medium simultaneously during siRNA transfection. To test the stability of cytoplasmic RNAs, 5 *μ*g/mL actinomycin D (ActD, Sigma-Aldrich, A9415) was added to the medium. For the cell viability test, the Cell Counting Kit-8 (CCK-8; Dojindo, CK04- 13) was used following the manufacturer’s protocol.

### Subcellular fractionation

Cytoplasmic fraction was obtained using the NE-PER Nuclear and Cytoplasmic Reagent Kit (Thermo Fisher Scientific, 78833). Cells were sequentially lysed with CER I and CER II buffers at 4°C for 10 min and 1 min, respectively. The supernatant (cytoplasmic fraction) was collected by centrifuging the lysates at 20,000 × *g* at 4°C for 5 min.

### RNA extraction and RT-qPCR

Total RNA was extracted from cells or fractionated lysates using TRIzol (Thermo Fisher Scientific, 15596018) or TRIzol LS (Thermo Fisher Scientific, 10296028), respectively. Following chloroform extraction and precipitation, DNase I (Takara, 2270B) was treated to eliminate the residual DNA, and purified RNA was reverse transcribed using RevertAid reverse transcriptase (Thermo Fisher Scientific, EP0441) or SuperScript IV reverse transcriptase (Thermo Fisher Scientific, 18090050) with random hexamer primers (Thermo Fisher Scientific, 48190-011). For lariat RNA-specific RT-qPCR, reverse transcription primers spanning the branch point were designed to target lariat RNAs. RT-qPCR was performed using the SensiFAST SYBR Lo-ROX Kit (Bioline, BIO-94005) and analyzed on QuantStudio 1 (Thermo Fisher Scientific). The primer sequences used in this study are listed in **Tables 3 and 4**.

### Western blotting

Cell lysates were prepared by sonication in lysis buffer (50 mM Tris-HCl pH 8.0, 100 mM KCl, 0.5% NP-40, 10% glycerol, and 1 mM DTT) supplemented with protease inhibitor cocktail (Thermo Fisher Scientific, 87785) and phosphatase inhibitor cocktail III (Sigma-Aldrich, P0044). 40-50 μg of proteins were separated on an SDS-PAGE gel and transferred to a polyvinylidene difluoride (PVDF) membrane using the Amersham semi-dry transfer system. For primary antibodies, anti-pPKR (Abcam, ab81303), anti-PKR (Cell Signaling Technology, 12297), anti-PKR (Santa Cruz Biotechnology, sc-6282), anti-peIF2α (Cell Signaling Technology, 9721), anti-eIF2α (Cell Signaling Technology, 5324), anti-DBR1 (Proteintech, 16019-1-AP), anti-MDA5 (Cell Signaling Technology, 5321), anti-RIG-I (Cell Signaling Technology, 3743), anti-ADAR1 (Cell Signaling Technology, 14175), anti-G3BP1 (Cell Signaling Technology, 17798), anti-PARP (Cell Signaling Technology, 9542), anti-Caspase-8 (Cell Signaling Technology, 9746), anti-cleaved Caspase-3 (Cell Signaling Technology, 9661), anti-Cyclin B1 (Cell Signaling Technology, 12231), anti-Lamin A/C (Santa Cruz Biotechnology, sc-376248), anti-p-Histone H3 (Cell Signaling

Technology, 3377), anti-Histone H3 (Cell Signaling Technology, 4499), anti-TUBB (Cell Signaling Technology, 86298), and anti-GAPDH (Santa Cruz Biotechnology, sc-32233) were used.

### J2 dot-blot assay

The 0.5 μL aliquots of RNAs diluted in nuclease-free water were loaded onto a positively charged nylon membrane (Amersham Hybond-N+, Cytiva, RPN1210B). The membrane was air-dried at RT for 45 min and blocked with 5% (w/v) skim milk diluted in TBST buffer (0.1% Tween 20). For the detection of dsRNAs, the membrane was incubated overnight at 4°C with anti-dsRNA J2 monoclonal antibody (SCICONS, 10010500) (diluted 1:500 in TBST containing 1% (w/v) skim milk). Following incubation, the membrane was washed 3 times with 0.1% TBST for 10 min each and was incubated for 1 h at RT with HRP-Goat Anti-Mouse IgG (H+L) Secondary Antibody (Jackson ImmunoResearch, 115-035-146) (1:3,000 diluted by TBST buffer containing 1% (w/v) skim milk). After washing the membrane 4 times with TBST, the signal was detected using SuperSignal^TM^ West Pico Plus chemiluminescent substrates (Thermo Fisher Scientific, 34580).

### Formaldehyde-mediated crosslinking and immunoprecipitation

To conjugate PKR antibody (Cell Signaling Technology, 12297) to beads, Pierce Protein A UltraLink Resin (Thermo Fisher Scientific, 53139) was incubated with the antibody in fCLIP lysis buffer (20 mM Tris-HCl pH 7.5, 15 mM NaCl, 10 mM EDTA, 0.5% NP-40, 0.1% Triton X-100, 0.1% sodium dodecyl-sulfate (SDS), and 0.1% sodium deoxycholate) with gentle rotation at 4°C for 3 h, after adjusting NaCl concentration to 150 mM. Harvested cells were fixed with 0.1% (w/v) filtered paraformaldehyde (Sigma-Aldrich, P6148) for 10 min, followed by quenching with glycine (final concentration of 250 mM) for 10 min. The crosslinked cells were lysed by incubation with fCLIP lysis buffer on ice for 10 min and then sonicated. Cell debris was removed by centrifugation after the concentration of NaCl in the lysate was adjusted to 150 mM as well. The lysate was then incubated with the PKR antibody-conjugated beads with gentle rotation at 4°C for 3 h. The beads were washed 4 times using fCLIP wash buffer (20 mM Tris-HCl pH 7.5, 150 mM NaCl, 10 mM EDTA, 0.1% NP-40, 0.1% SDS, 0.1% Triton X-100, and 0.1% sodium deoxycholate). The PKR-RNA complex was eluted from the beads by incubation in the elution buffer (200 mM Tris-HCl pH 7.4, 100 mM NaCl, 20 mM EDTA, 2% SDS, and 7 M Urea) at 25°C for 3 h. To de-crosslink RNAs from PKR, 2 mg/ml proteinase K (Roche, 3115879001) was added to the eluate and incubated at 65°C overnight. After de-crosslinking, acid-phenol:chloroform pH 4.5 was added to purify RNAs. For J2 fCLIP, the same protocol was followed using the J2 antibody (SCICONS, 10010500).

### Immunofluorescence

Cells cultured in a 0.1% (w/v) gelatin-coated confocal dish were washed once with PBS and fixed with 4% (w/v) paraformaldehyde at RT for 10 min. For RNase R and RNase A treatment, prior to paraformaldehyde fixation, cells were permeabilized with 0.02% (v/v) Triton X-100 in Tris buffer (20 mM Tris-HCl pH 7.4, 5 mM MgCl_2_, 0.5 mM EDTA, 25% Glycerol, and 100 mM KCl) at RT for 3 min, followed by incubation with RNase R (20 U/mL, Lucigen, RNR07250) or RNase A (1 mg/mL, Thermo Fisher Scientific, EN0531) at 37°C for 10 min. The fixed cells were permeabilized with 0.5% (v/v) Triton X-100 in PBS for 10 min and then blocked with 1% (w/v) BSA at RT for 1 h. Subsequently, cells were incubated with primary antibodies diluted in 1% BSA at RT for 2 h. Cells were then rinsed twice and washed three times with 0.1% (v/v) Tween 20 in PBS (PBST) and incubated with Alexa fluorophore labeled secondary antibodies at RT for 45 min. To counterstain the nuclei, 450 nM of 4**′**,6-diamidino-2-phenylindole dihydrochloride (DAPI) (Thermo Fisher Scientific, D1306) was added to the secondary antibody solution.

For total RNA staining, cells were fixed 4% (w/v) paraformaldehyde at RT for 10 min. Following three PBS washes, cells were permeabilized with 0.5% (v/v) Triton X-100 in PBS at RT for 10 min. Subsequently, cells were incubated in SYTO® RNASelect Green Fluorescent Stain (Thermo Fisher Scientific, S32703) staining solution (1 μM in PBS) at RT for 20 min, followed by three additional PBS washes, with 450 nM of DAPI included in the staining solution to counterstain the nuclei. All fluorescence images were acquired using the Zeiss LSM780 confocal microscope.

### Immunohistochemical Staining

Paraffin-embedded tumor tissues were sectioned at 4 μm thickness and processed for immunohistochemistry using the Bond-RXm automated staining system (Leica Biosystems). The following primary antibodies were applied: anti-DBR1 (Proteintech, 16019-1-AP, 1:500), anti-phospho-PKR (Thermo Fisher Scientific, 44-668G, 1:100), and anti-PKR (Santa Cruz Biotechnology, sc-6282, 1:200). Staining intensity was graded using an intensity score (IS) on a 0–3 scale: 0 (negative), 1 (weak), 2 (moderate), and 3 (strong). The proportion score (PS) was defined as the percentage of tumor cells exhibiting each staining intensity, with the total summing to 100%. To quantitatively assess expression levels, the H-score was calculated for each sample using the formula: H-score = (IS_0_ × PS_0_) + (IS_1_ × PS_1_) + (IS_2_ × PS_2_) + (IS_3_ × PS_3_), yielding a score range of 0 to 300. All stained slides were independently evaluated by two board-certified breast pathologists (S.Y.P. and H.S.R.), and any discrepancies were resolved by consensus review.

### RNA-fluorescent in situ hybridization (RNA-FISH)

RNA-FISH probes were prepared using the MEGAscript T7 Transcription Kit (Thermo Fisher Scientific, AM1334) and DIG RNA Labeling Mix (Roche, 11277073910). DNA templates for in vitro transcription (IVT) were amplified using KOD-Plus-Neo (TOYOBO, KOD-401) from cDNA with a reverse primer containing the T7 promoter sequences at the 5′ end. IVT was performed using a DIG labeling mix at 37°C for 6 h, subsequently treated with Turbo DNase I digestion at 37°C for 15 min and purified using Monarch RNA Cleanup Kit (NEB, T2040). The purified RNA probe was fragmented in the hydrolysis buffer (40 mM NaHCO_3_ and 60 mM Na_2_CO_3_ pH 10) at 60°C for 40 min, and then quenched with the stop buffer (0.2 M NaOAc pH 6) on ice. The RNA probe was then diluted in the hybridization buffer (50% formamide, 10% dextran sulfate, 0.1% SDS, 300 ng/mL salmon sperm DNA, 2X SSC, and 2 mM vanadyl ribonucleoside complex).

Cells grown on a confocal dish were fixed with 4% (w/v) paraformaldehyde at RT for 10 min and permeabilized in 70% ethanol (in 2X SSC) overnight at 4°C. The cells were rehydrated by successive washing with 50% ethanol (in 2X SSC) and 25% ethanol (in 2X SSC) at RT for 3 min each. Cells were incubated in 50% formamide (in 2X SSC) at 40°C for 2 h and hybridized overnight at 40°C with a denatured RNA probe. Hybridized cells were stringently washed twice with 50% formamide (in 2X SSC) and 50% formamide (in 0.1X SSC) at 40°C for 30 min each. The cells were blocked in the blocking buffer (0.2% BSA, 8% formamide, 2X SSC) at RT for 1 h. Cells were incubated with anti-DIG antibody (Roche, 11333089001) diluted in the blocking buffer at RT for 2 h, followed by labeling using Alexa Fluor-conjugated secondary antibody. Nuclei were counterstained with DAPI, and imaging was performed with a Zeiss LSM780 confocal microscope.

### Protein expression and purification

12 μg of expression plasmid constructs (pcDNA3.1-DBR1-FLAG and pCK-FLAG-PKR) for FLAG- tagged proteins (DBR1 and PKR) were individually transfected into 10 mL of HEK-293T cell culture (6.0 x 10^5^ cells/mL) using FuGENE HD transfection reagent (Promega, E2311) following the manufacturer’s protocol. The cells were harvested 48 h after transfection, and the pellets were suspended in 500 μL of lysis buffer (50 mM Tris-HCl pH 8.0, 100 mM KCl, 0.5% NP-40, 10% glycerol, and 1 mM DTT) containing protease inhibitor cocktail. The lysate was sonicated, followed by centrifugation (20,000 × *g*, 10 min). The supernatant was aliquoted and incubated with 40 μL of anti-FLAG M2 affinity gel (Sigma-Aldrich, A2220) at 4°C for 4 h with gentle rotation. After incubation, the affinity gel was washed 4 times with 500 μL of FLAG buffer (50 mM Tris-HCl pH 7.4 and 150 mM NaCl). The remaining buffer was fully removed, and 60 μL of FLAG buffer containing 0.5 mg/mL 3X FLAG peptide (Thermo Fisher Scientific, A36805) was directly added to the affinity gel. The elution mixture was incubated in ThermoMixer C (Sigma-Aldrich) (1,300 rpm, 4°C for 30 min) and centrifuged (6,000 × *g*, 1 min). The supernatant was collected and stored at –80°C.

### DBR1 mutagenesis and overexpression

To generate the DBR1 catalytic mutant (DBR1 H85N), the DBR1 WT plasmid (GenScript, OHu11162) was amplified with the primer with point mutations using a high-fidelity nPFU-Forte (Enzynomics, P410) according to the manufacturer’s instructions. Following PCR, DpnI was used to degrade the methylated DNA template, and unmethylated PCR products were transformed into *E. coli* DH5α competent cells. The point mutations were verified through the Sanger sequencing.

To establish MDA-MB-231 cells overexpressing either DBR1 WT or H85N, the Cas9 sequence in lentiCas9-Blast (Addgene, 52962) was replaced with the coding sequence of DBR1 WT or H85N, generating lentiDBR1-WT-Blast and lentiDBR1-H85N-Blast. Lentivirus particles were generated in HEK-293T cells by co-transfecting lentiDBR1-WT-Blast or lentiDBR1-H85N-Blast with psPAX2 (Addgene, 12260) and pMD2.G (Addgene, 12259) supplemented by 1% (v/v) BSA. Media containing lentivirus particles were harvested after 60 h of transfection and transduced into MDA-MB-231 cells. Two days after transduction, blasticidin (Gibco, A1113903) was added to the media for selection.

### In vitro PKR binding assay

Total RNAs extracted from DBR1-deficient MDA-MB-231 cells were treated with RNase R and recovered as described in a previous study.^60^ RNase R-treated RNAs were heated at 65°C for 5 min in RNA-folding buffer (10 mM HEPES and 10 mM MgCl_2_) and slowly cooled down to RT. 3 μg of folded RNAs were incubated with 1.5 μg of recombinant PKR (Origene, TP326606) at 4°C for 2 h in 0.2 mL Binding buffer (50 mM HEPES at pH 7.0, 150 mM NaCl, 10 mM MgCl_2_, 0.1 mM DTT) supplemented with protease inhibitor cocktail and recombinant RNase I inhibitor (Takara, 2313A). After binding, the mixture was added to the PKR antibody-conjugated beads and incubated at 4°C for 3 h with gentle rotation. The beads were washed 4 times using 0.5 mL of Binding buffer containing 0.1% NP-40. After washing, PKR-binding RNAs were recovered with TRIzol.

### In vitro PKR activation assay

In vitro activation assay of PKR was performed as described with some modifications. ^61^ Purified recombinant PKR was first dephosphorylated by treating it with λ-PPase (NEB, P0753). Subsequently, λ-PPase was inactivated by treatment with freshly prepared sodium orthovanadate (NEB, P0758). Next, 300 ng of dephosphorylated PKR and different RNAs were incubated in 20 *μ*L of in vitro activation buffer (20 mM HEPES pH 7.5, 4 mM MgCl_2_, 100 mM KCl, and 1 mM ATP) at 30°C for 30 min. Reactions were stopped by adding an SDS loading buffer and analyzed on a 10% SDS-PAGE gel. Phosphorylated PKR and total PKR amounts were assessed by western blotting with antibodies against pPKR (Abcam, ab81303) and PKR (Cell Signaling Technology, 12297), respectively.

### In vitro debranching assay

In vitro debranching assay was performed with a total reaction volume of 50 μL with 300 ng of purified DBR1-FLAG protein and 2.5 μg of RNAs in the debranching buffer (40 mM Tris-HCl pH 7.0, 80 mM KCl, 6 mM MgCl_2_, 8% Glycerol) supplemented with protease inhibitor cocktail and recombinant RNase I inhibitor. The debranching mixture was incubated at 37°C for 1 h. After incubation, RNAs were recovered with acid phenol extraction and ethanol precipitation.

### In vitro synthesis of lariat-mimic circular intron RNAs

Lariat-mimic circular intron RNAs were synthesized using the Complete self-splicing Intron for RNA Circularization (CIRC method).^37^ The CIRC template comprises T7 promoter and an *Anabaena* group I intron followed by a minimal 2 nt 3′ exon (AA), the target intron sequence (payload sequence), and a minimal 5′ exon (TCT). Genomic sequences spanning from the 5′ splice site to the branch point of *DKK1* intron 1, *CANX* intron 5, and *PTBP1* intron 13 were cloned into the CIRC template and verified by Sanger sequencing. The sequences of all CIRC constructs are provided in **Table 5**.

For *DKK1* intron 1 and *CANX* intron 5, IVT was performed using the MEGAscript T7 kit at 37 °C for 4 h, since both payloads yielded abundant circular product directly from the precursor under these conditions. After DNase I treatment, the RNA was purified using the Monarch RNA Cleanup Kit. Circular intron RNAs were then resolved on a 4% urea-PAGE gel, and the circular band was extracted and recovered, followed by RNase R treatment to remove residual linear RNA contaminants.

For *PTBP1* intron *13*, which is larger and prone to substantial byproduct circularization occurring during IVT, cyclic GMP (Sigma-Aldrich, G7504) was included in the IVT reaction (GTP : cGMP = 7.5 mM : 15 mM) to suppress group I intron activity and enrich for the precursor. Following Monarch cleanup purification, the precursor was resolved on a 4% urea-PAGE gel, and the precursor band was excised to remove byproducts. The purified precursor was then circularized by refolding in the Circularization buffer (50 mM HEPES pH 7.5 and 6 mM MgCl₂) at 55°C for 1 h. The circular intron RNA was resolved on a 0.8% agarose gel, and circular band was excised and recovered by acid-phenol extraction: the gel slice was soaked in 5 volumes of soaking buffer (0.3 M NaOAc and 1 mM EDTA), frozen at −80°C for 30 min, thawed at 65°C for 10 min with intermittent vortexing, and extracted with an equal volume of acid phenol:chloroform (pH 4.5). The aqueous phase was ethanol-precipitated at −80°C for 1 h, washed three times with 75% ethanol, and resuspended in nuclease-free water. Purified RNAs were treated with RNase R to eliminate residual linear RNA species. Gel-extracted, RNase R-treated circular intron RNAs were used for in vitro PKR activation assay and transfected into cells using Lipofectamine 3000.

### RNA-seq library preparation and alignment to the genome

Paired-end total RNA-seq and in vitro PKR binding RNA-seq were constructed using the TruSeq Stranded Total RNA with Ribo-zero H/M/R/ Gold kit (Illumina) and sequenced using the NovaSeq 6000 platform (Illumina) or NovaSeqX (Illumina). For the in vitro debranching assay RNA-seq and PKR fCLIP-seq, SMARTer Stranded Total RNA-seq Kit (Takara) was used and sequenced using NovaSeqX (Illumina). Raw paired-end sequencing reads were mapped to the reference human genome (build hg38) using HISAT2 v2.1.0^62^ with default parameters, except with the option “--rna-strandness”. Uniquely and concordantly mapped reads were selected using samtools^63^ and then were used for further analyses. The distribution of in vitro PKR binding RNA-seq reads across genomic features was analyzed using RSeQC v4.0.0^64^ using the read_distribution.py module.

### Identification of intronic genomic regions and intronic transposable elements

A comprehensive list of human introns was generated using the latest transcriptomic annotation (GENCODE v34)^65^. Briefly, intronic regions were defined as the spanning intervals between two adjacent exons across all protein-coding genes. Only introns with lengths shorter than 30,000 bp were used for the analysis. To identify transposable elements residing in intronic regions, the genomic location information was obtained from the most recent version of Repeatmasker^66^ for hg38. Specifically, sense and antisense *Alu* elements residing in intronic regions were identified based on their orientation relative to the corresponding introns. Intron RNAs containing at least one sense and at least one antisense *Alu* were classified as IR*Alus*-containing intron RNAs. Intron RNAs containing two or more sense *Alus* only or two or more antisense *Alus* only without their complementary pairs were classified as DR*Alus*-containing intron RNAs.

### Annotation of RNA-seq alignments and data processing

To assign and summarize genomic features, the featureCounts function of the Rsubread v2.8.2^67^ R package was used with transcriptome information GENCODE v34 or Repeatmasker pre-masked for hg38. For GENCODE, the transcriptome information of only protein-coding transcripts was included. To calculate the in vitro PKR enrichment ratio, raw read counts normalized to RPM were used. TEtranscripts^68^ was employed to quantify the read count and fold change of each TE sub-family.

For GSEA, the clusterProfiler v4.2.2^69^ and DOSE v.3.20.1^70^ R packages were used. Hallmark gene sets were obtained from the Molecular Signatures Database (MSigDB v7.4)^71^. For lariat read analysis, branchpoint-traversing lariat reads were identified using an iterative inverted alignment method described in the previous study^72^. Forward alignment was performed using the STAR RNA-seq aligner^73^, and the lariat read recovery rate was calculated as the number of recovered lariat reads per million mapped reads to hg19.

### A-to-I editing score

REDItools2^74^ was used to identify and quantify RNA editing sites from total RNA-seq of MDA-MB- 231 cells transfected with siDBR1 for 6 days, using default parameters while considering strandness specificity. To confirm the A-to-I editing, intron RNAs with Log_2_(siDBR1 RPM/siLuc RPM) greater than 2 in total RNA-seq data were analyzed. A-to-G mismatch sites (read depth ≥ 5) detected in the sequencing reads aligned to intronic regions were identified as intronic A-to-I editing sites.

The A-to-I editing score for each intron RNA was defined as the sum of the A-to-I editing rate normalized to its length: 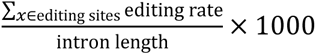. A-to-I editing sites with an editing rate between 0.05 and 0.95 were counted to exclude potential SNPs or random errors in sequencing. For intronic C-to-U editing, C-to-T mismatch sites in the sequencing reads were analyzed following the same criteria applied to A-to-I editing.

### Quantification and statistical analysis

Statistical analyses and data visualization were performed using RStudio (R version 4.3.0). The number of biological replicates and type of statistical tests used are indicated in the figure legends.

## Data availability

Total RNA-seq following subcellular fractionation, in vitro PKR binding RNA-seq, and PKR fCLIP- seq data have been deposited in GEO under accession number: GSE287454. Original western blot images, gel images, and microscopy data have been deposited at Mendeley (DOI: 10.17632/h59mmzbdc4.1).

