## Extended Data Figs and Tables for "Nuclear retention and RNA editing safeguard self-tolerance to immunogenic intron lariats"

### Extended Data Figs. 1-8 and Tables 1-5

#### Extended Data Fig. 1. Effects of DBR1 depletion on intron lariat RNAs and cellular dsRNAs

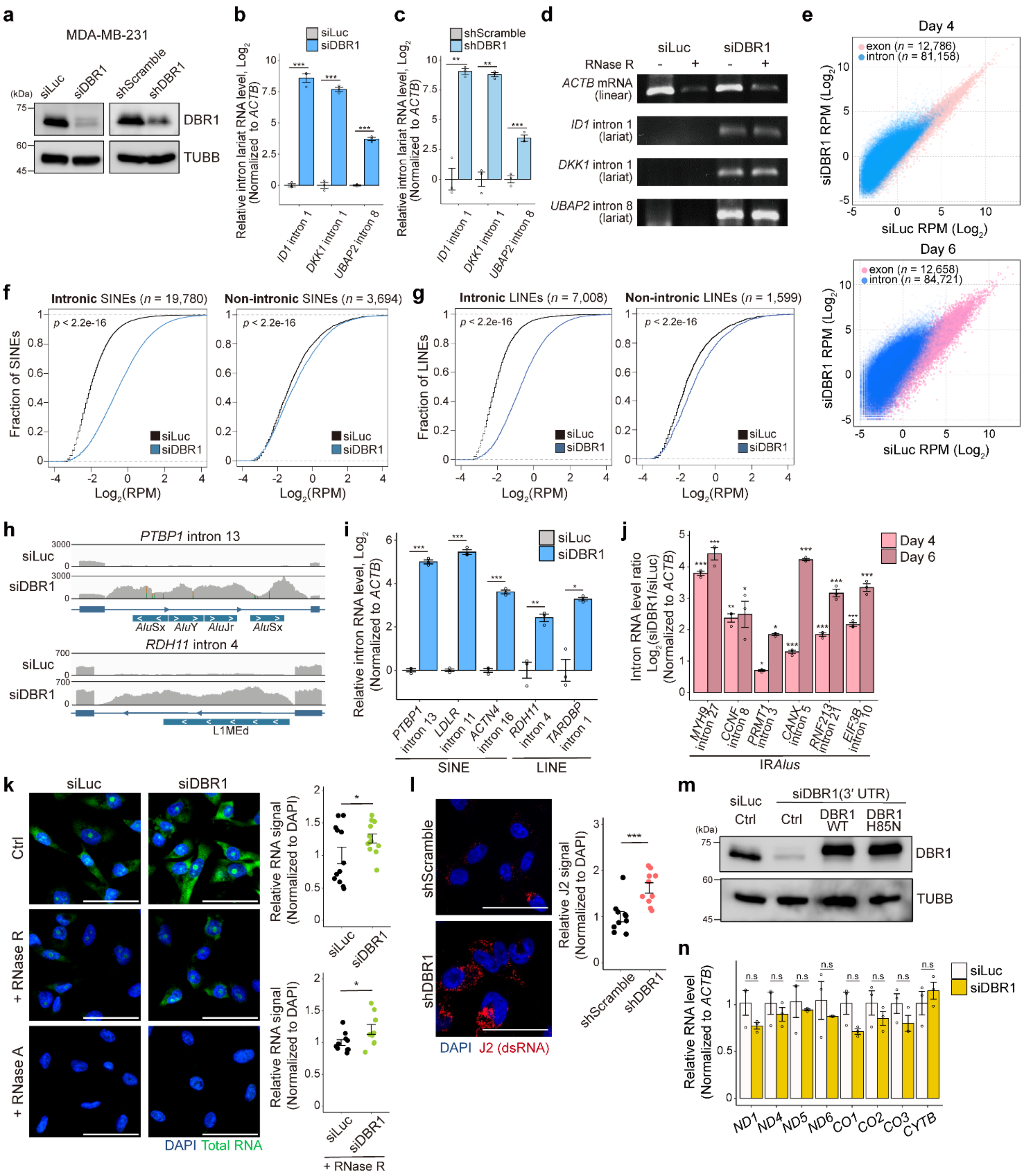

**a.** KD efficiency of DBR1 in MDA-MB-231 transfected with siDBR1 or transduced with shDBR1, analyzed by western blotting (WB).

**b, c.** RT-qPCR analysis of selected intron lariat RNAs 6 days after siDBR1 transfection (**b**, $n$ = 3) or shDBR1 transduction (**c**, $n$ = 3) to MDA-MB-231 using lariat-specific divergent primers.

**d.** RT-PCR of intron lariat RNAs, using lariat-specific primers, after RNase R treatment on total RNAs extracted from MDA-MB-231 transfected with siDBR1 for 6 days.

**e.** Scatter plots comparing mean exon and intron RPMs for MDA-MB-231 transfected with siRNA for 4 (left) and 6 (right) days. Exons are shown in pink, and introns are shown in blue.

**f.** Cumulative distribution curves of mean RPMs for intronic (left, $n$ = 19,780) and non-intronic SINEs (right, $n$ = 3,694) in MDA-MB-231 transfected with siRNAs for 6 days.

**g.** Cumulative distribution curves of mean RPMs for intronic (left, $n$ = 7,008) and non-intronic LINEs (right, $n$ = 1,599) in MDA-MB-231 transfected with siRNAs for 6 days.

**h.** IGV snapshots of total RNA-seq reads at *PTBP1* intron 13 and *RDH11* intron 4 from MDA-MB-231 transfected with siRNAs for 6 days.

**i.** RT-qPCR analysis of selected intron lariat RNAs containing either SINE or LINE 4 days after siDBR1 transfection to MDA-MB-231 ($n$ = 3).

**j.** RT-qPCR analysis of intron lariat RNAs containing IR*Alus* 4 or 6 days after siDBR1 transfection to MDA-MB-231 ($n$ = 3).

**k.** Comparison of total RNA levels (measured by SYTO RNA staining) following RNase R or RNase A treatment in MDA-MB-231 transfected with siRNAs for 6 days. Quantification is shown on the right ($n$ = 12 for control samples; $n$ = 10 for RNase R-treated samples).

**l.** Immunocytochemistry of J2 dsRNA signals for MDA-MB-231 6 days after shRNA transduction. Quantification is shown on the right ($n$ = 9).

**m.** DBR1 expression after transfection of siDBR1 targeting the 3′ UTR of *DBR1* mRNA and overexpression of DBR1 WT or H85N, analyzed by WB.

**n.** RT-qPCR analysis of mtRNAs 6 days after siDBR1 transfection to MDA-MB-231 ($n$ = 3).

All error bars denote s.e.m. All scale bars indicate 50 μm. $P$-values were calculated using either the one-sided Student's t-test (b, c, i, j, k, l, n) or the two-sided Mann-Whitney test (f, g); $^{*}p < 0.05$, $^{**}p < 0.01$, $^{***}p < 0.001$.

**Extended Data Fig. 2. DBR1 depletion-induced PKR activation**

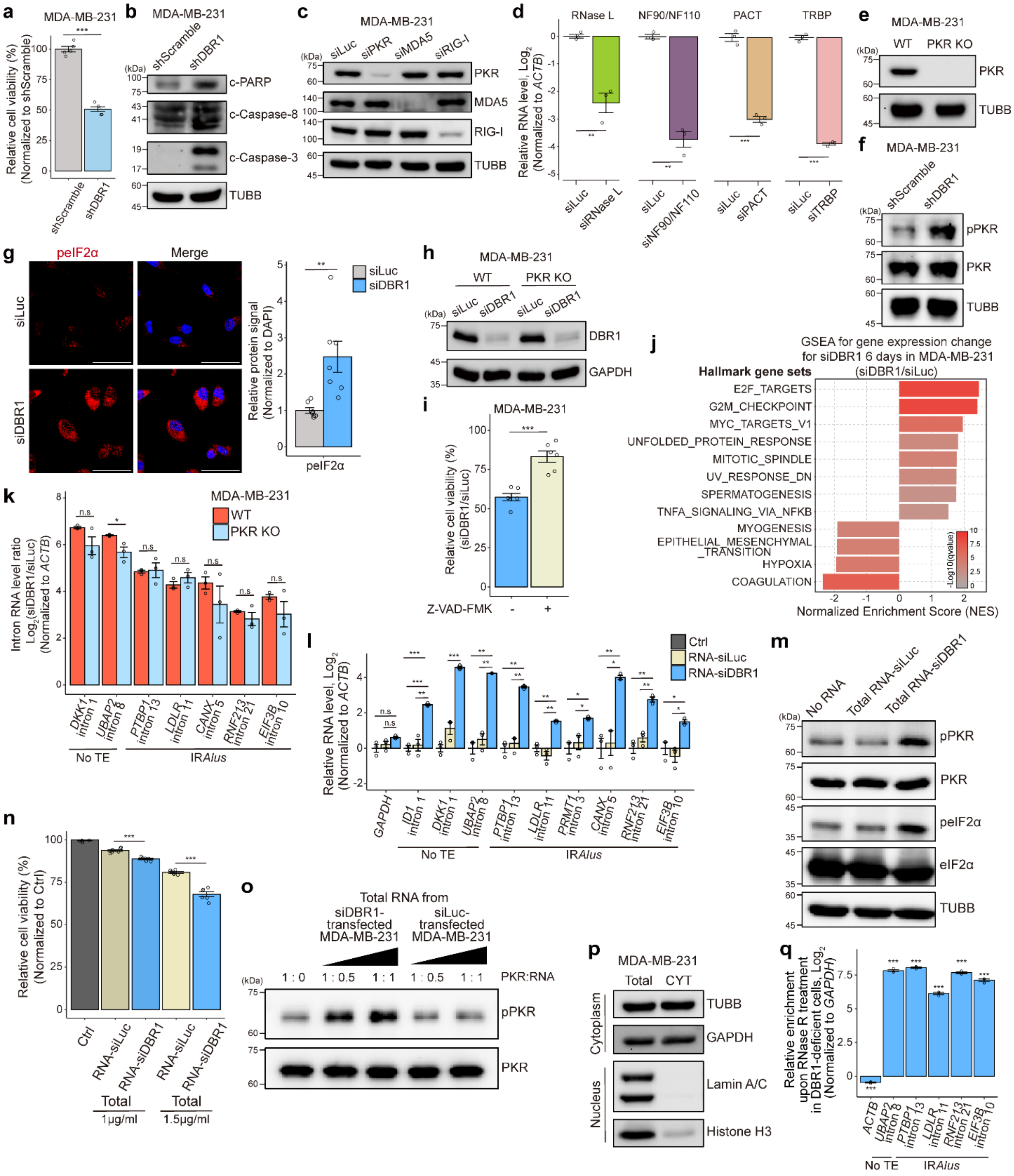

**a, b.** Cell viability (**a**, $n$ = 5) and WB of apoptotic markers (b) 6 days after shDBR1 transduction to MDA-MB-231.

**c.** KD efficiency of PKR, MDA5, and RIG-I in MDA-MB-231 following transfection with their respective siRNAs, analyzed by WB.

**d.** KD efficiency of RNase L, NF90/NF110, PACT, and TRBP in MDA-MB-231 following transfection with their respective siRNAs, analyzed by RT-qPCR.

**e.** Confirmation of PKR depletion in MDA-MB-231 PKR KO cells, analyzed by WB.

**f.** Phosphorylation of PKR 6 days after shDBR1 transduction to MDA-MB-231, analyzed by WB.

**g.** Phosphorylation of eIF2α 4 days after siDBR1 transfection to MDA-MB-231, analyzed by immunocytochemistry. Quantification is shown on the right ($n$ = 7). Scale bar indicates 50 μm.

**h.** KD efficiency of DBR1 after siDBR1 transfection to MDA-MB-231 WT and PKR KO, analyzed by WB.

**i.** Effect of the pan-caspase inhibitor on cell viability ($n$ = 6) in DBR1-deficient MDA-MB-231.

**j.** Significantly enriched or depleted Hallmark gene sets identified by GSEA for relative expression in MDA-MB-231 transfected with siDBR1 for 6 days.

**k.** RT-qPCR analysis of selected intron RNAs after siDBR1 transfection to MDA-MB-231 WT and PKR KO cells ($n$ = 3).

**l-n.** RT-qPCR analysis of intron RNAs (**l**, $n$ = 3), WB of PKR and eIF2α phosphorylation (**m**), and cell viability (**n**, $n$ = 6) 24 h after re-transfection of total RNA-siDBR1 or RNA-siLuc to MDA-MB-231.

**o.** In vitro PKR activation assay using recombinant PKR and total RNA-siDBR1 or RNA-siLuc. The ratio indicates the mass ratio of RNA to PKR.

**p.** Successful fractionation of cytoplasm (TUBB and GAPDH: Cytoplasm / Lamin A/C and Histone H3: Nucleus), analyzed by WB of marker proteins.

**q.** Relative enrichment of intron RNAs following RNase R digestion of total RNA isolated from siDBR1-transfected MDA-MB-231 ($n$ = 3).

All error bars denote s.e.m. $P$-values were calculated using the one-sided Student's t-test (a, d, g, i, k, l, n, q); $^{*}p < 0.05$, $^{**}p < 0.01$, $^{***}p < 0.001$, and n.s: non-significant.

#### Extended Data Fig. 3. Characterization of intron lariat RNA-PKR interactions

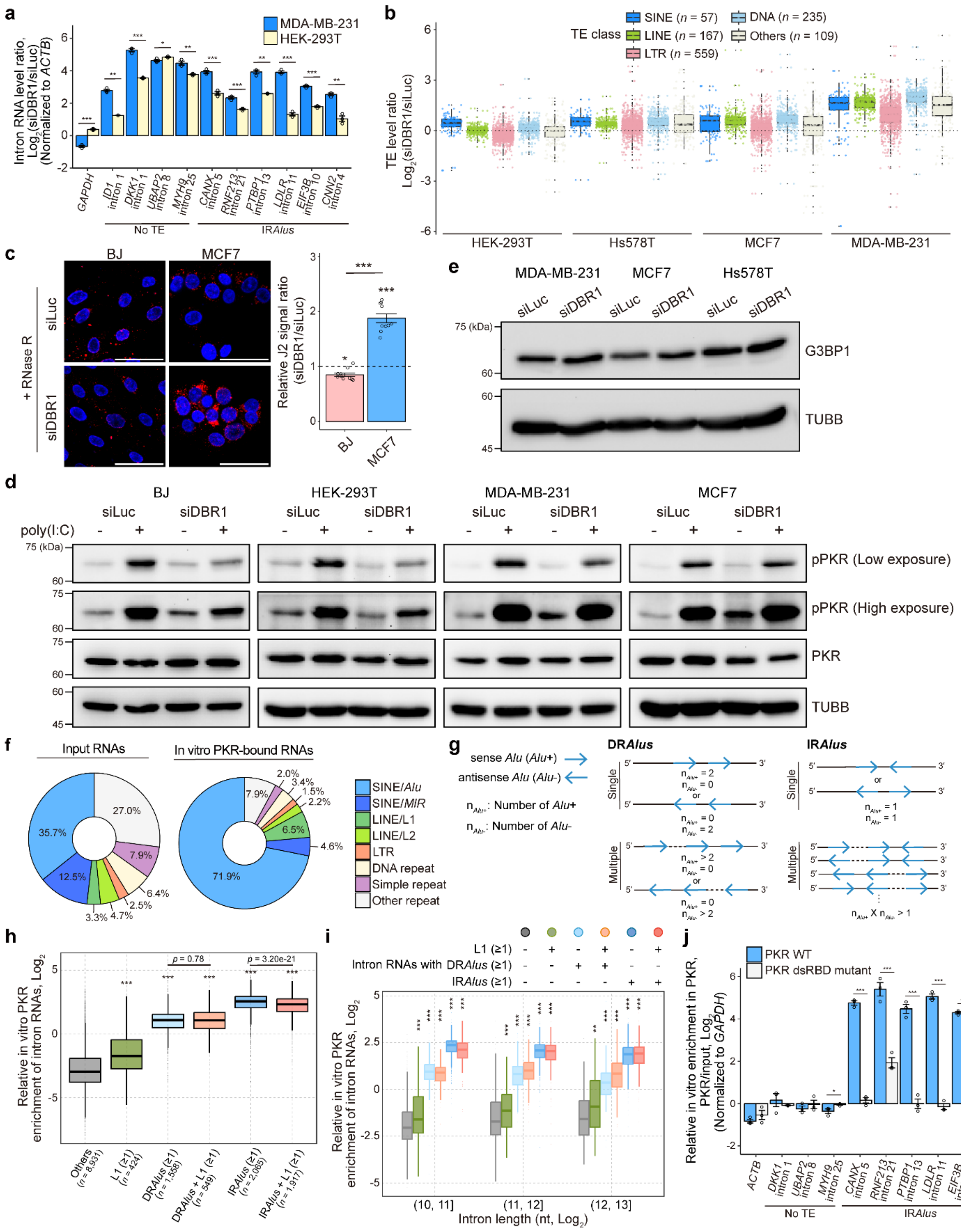

**a.** Relative levels of selected intron RNAs 6 days after siDBR1 transfection to MDA-MB-231 and HEK-293T (*n* = 3), analyzed by RT-qPCR.

**b.** TE expression changes 6 days after siDBR1 transfection to HEK-293T, Hs578T, MCF7, and MDA-MB-231. Each point denotes a specific TE sub-family.

**c.** Comparison of J2 dsRNA signal following RNase R treatment 6 days after siRNA transfection to BJ and MCF7. Quantification is shown on the right (*n* = 10). Scale bar indicates 50 μm.

**d.** WB of PKR phosphorylation in BJ, HEK-293T, MDA-MB-231, and MCF7 6 days after siDBR1 transfection, followed by poly(I:C) (500 ng/mL) transfection for 2 h.

**e.** WB of G3BP1 expression 4 days after siDBR1 transfection to MDA-MB-231, MCF7, and Hs578T.

**f.** Relative proportion of in vitro PKR binding-seq reads (from siDBR1-transfected MDA-MB-231) for various TE types.

**g.** Definition of *Alu* association (DR*Alus* and IR*Alus*) depending on the number and orientation of *Alu* elements in a single intron. *Alu*+ denotes sense *Alu,* and *Alu*- denotes antisense *Alu*.

**h.** Impact of co-presence of L1 in DR*Alus* or IR*Alus*-containing intron RNAs on their relative in vitro PKR enrichments.

**i.** Impact of intron length on relative in vitro PKR enrichment of intron RNAs containing DR*Alus,* IR*Alus,* or L1. Intron RNAs lacking all *MIR*, *Alu*, L1, and L2 elements are referred to as “Others”.

**j.** RT-qPCR analysis of intron RNAs following in vitro PKR binding assay using purified PKR WT and PKR dsRBD mutant (PKR S59A K60A T149A K150A), normalized to input (*n* = 3).

All error bars denote s.e.m. *P*-values were calculated using either the one-sided Student’s t-test (a, c, j) or the two-sided Wilcoxon signed-rank test (h, i); *$p < 0.05$, **$p < 0.01$, ***$p < 0.001$.

#### Extended Data Fig. 4. In vitro synthesis of lariat-mimic circular intron RNAs

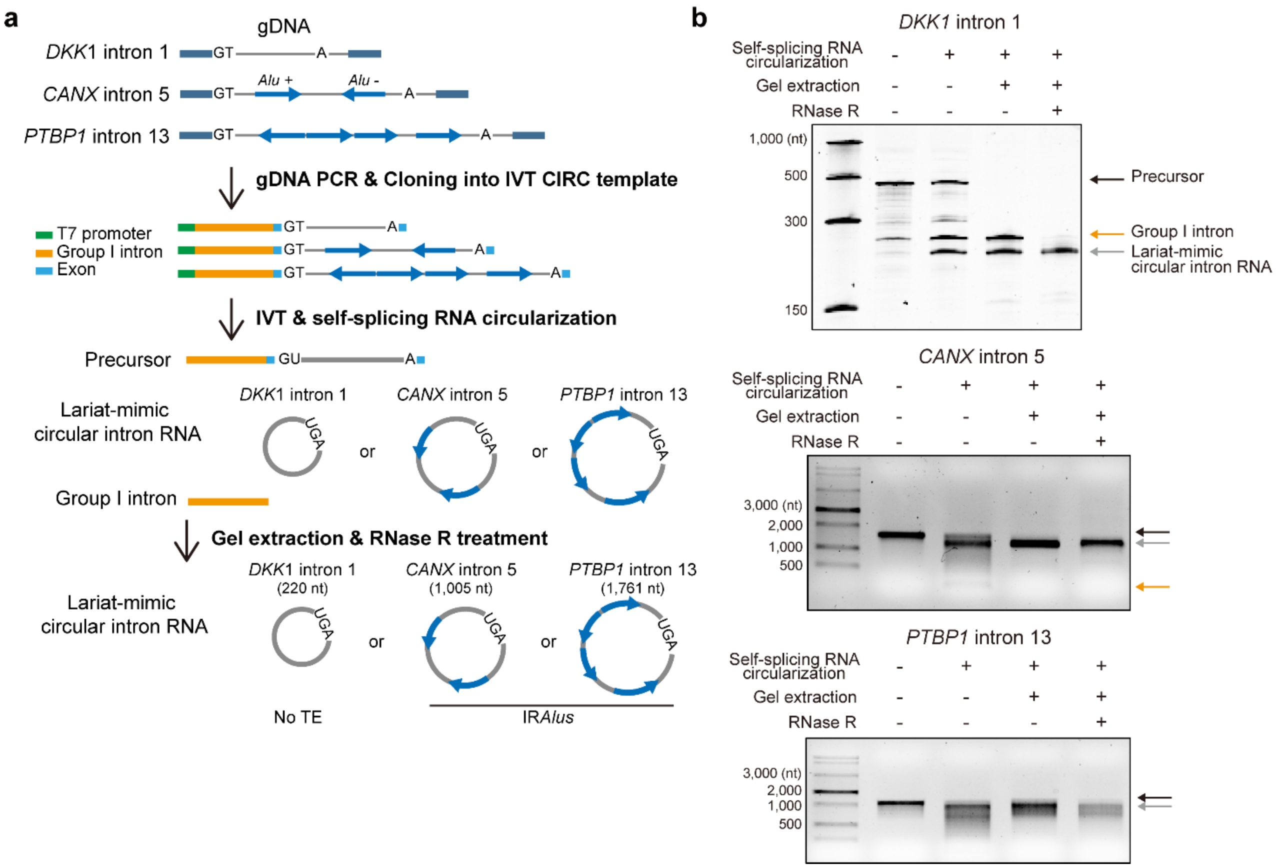


**a.** Experimental scheme for the preparation of lariat-mimic circular intron RNAs. Sequences of *DKK1* intron 1, *CANX* intron 5, and *PTBP1* intron 13 were PCR-amplified from MDA-MB-231 genomic DNA (gDNA) and cloned into IVT templates. Circular intron RNAs were generated by in vitro transcription (IVT) followed by group I intron-mediated self-splicing circularization, purified by gel extraction, and treated with RNase R to remove residual linear RNAs.

**b.** Quality assessment of in vitro synthesized circular intron RNAs, analyzed by 6% urea-PAGE gel for *DKK1* intron 1 and 1% agarose gel for *CANX* intron 5 and *PTBP1* intron 13.

#### Extended Data Fig. 5. Effects of lariat structure on intron RNA-PKR interactions

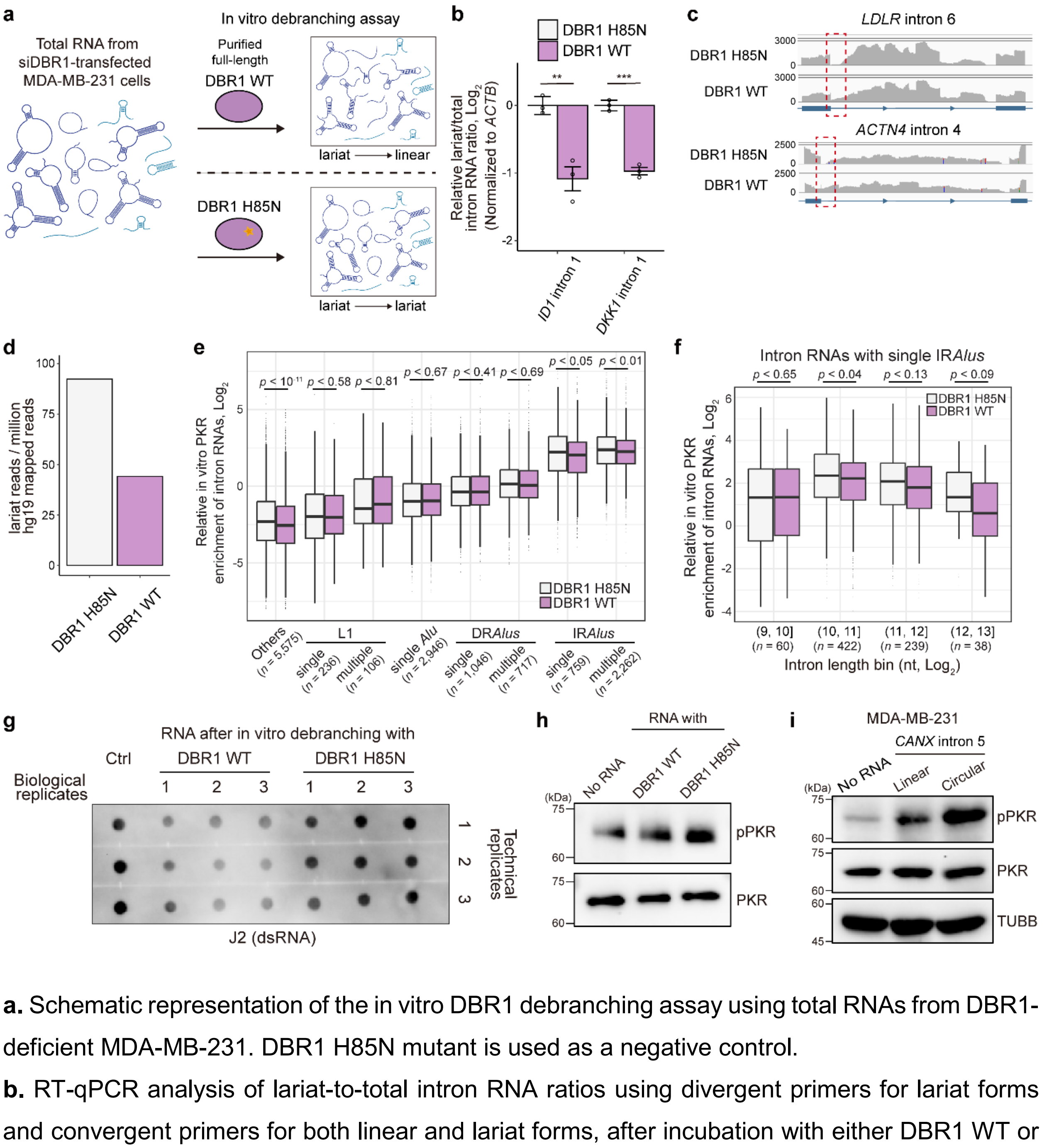


**a.** Schematic representation of the in vitro DBR1 debranching assay using total RNAs from DBR1-deficient MDA-MB-231. DBR1 H85N mutant is used as a negative control.

**b.** RT-qPCR analysis of lariat-to-total intron RNA ratios using divergent primers for lariat forms and convergent primers for both linear and lariat forms, after incubation with either DBR1 WT or H85N ($n$ = 3).

**c.** IGV snapshots of RNA-seq reads mapped at *LDLR* intron 6 and *ACTN4* intron 4 after in vitro debranching assay with DBR1 WT or H85N.

**d.** Lariat read counts from total RNA-seq after in vitro debranching assay with DBR1 WT or H85N, normalized against mapped read counts to hg19.

**e.** Impact of the lariat structure of intron RNAs on relative in vitro PKR enrichment, depending on the type and the number of L1 and *Alu* elements. Intron RNAs lacking all *MIR*, *Alu*, L1, and L2 elements are referred to as “Others”.

**f.** Impact of the lariat structure of intron RNAs with single IR*Alus* on relative in vitro PKR enrichment depending on the intron length.

**g.** J2 dot-blot assay of total RNA isolated from DBR1-deficient MDA-MB-231 after in vitro debranching. Three biological replicates with technical triplicates are shown. ‘Ctrl’ denotes input RNA used for the in vitro debranching assay.

**h.** In vitro PKR activation assay using recombinant PKR and total RNA isolated from DBR1-deficient MDA-MB-231 following in vitro debranching assay.

**i.** WB of PKR and eIF2α phosphorylation in MDA-MB-231 cells transfected with linear or circular *CANX* intron 5 RNA (500 ng/mL), assessed at 12 h post-transfection.

All error bars denote s.e.m. *P*-values were calculated using either the one-sided Student’s t-test (b) or the two-sided Wilcoxon signed-rank test (e, f); $^{**}p < 0.01$ and $^{***}p < 0.001$.

**Extended Data Fig. 6. Cytosolic release and subsequent PKR activation by intron lariat RNAs**

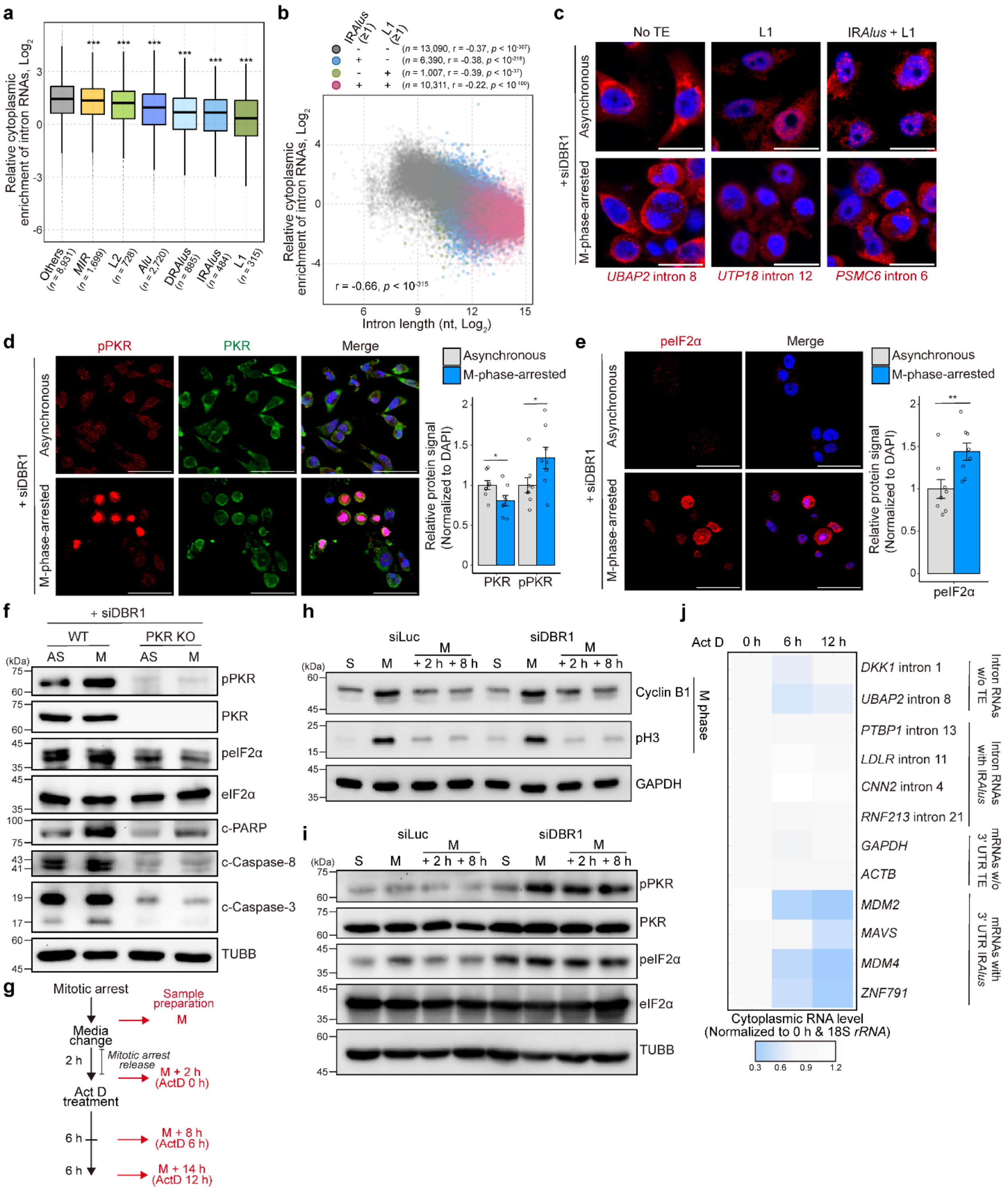

**a.** Relative cytoplasmic enrichment of intron RNAs containing either a single TE (*MIR*, *Alu*, L1, or L2), DR*Alus,* or IR*Alus*. Intron RNAs lacking all *MIR*, *Alu*, L1, and L2 elements are referred to as "Others".

**b.** Impact of intron length on relative cytoplasmic enrichment (cytoplasm/total RPM ratio) of intron RNAs containing either IR*Alus* or L1.

**c.** RNA-FISH analysis of selected intron RNAs upon M-phase arrest in DBR1-deficient MDA-MB-231. Scale bar indicates 20 μm.

**d, e.** Phosphorylation of PKR (**d**) and eIF2α (**e**) upon M-phase arrest in DBR1-deficient MDA-MB-231, analyzed by immunocytochemistry. Quantification is shown on the right (PKR, $n = 8$; eIF2α, $n = 8$). Scale bar indicates 50 μm.

**f.** WB of eIF2α phosphorylation and induction of apoptotic markers upon M-phase-arrest in DBR1-deficient MDA-MB-231 WT and PKR KO.

**g.** Experimental timeline for sample preparation after ActD treatment following release from mitotic arrest.

**h.** Confirmation of successful release from mitotic arrest by WB of M-phase marker proteins, Cyclin B1 and phosphorylated Histone H3 (pH3).

**i.** WB of PKR and eIF2α phosphorylation after release from mitotic arrest in MDA-MB-231.

**j.** Heatmap showing the relative cytoplasmic RNA level upon ActD treatment following release from mitotic arrest in DBR1-deficient MDA-MB-231 ($n = 3$), analyzed by RT-qPCR. Each square represents the mean RNA levels (normalized to ActD 0 h) across three biological replicates.

All error bars denote s.e.m. *P*-values were calculated using either the two-sided Wilcoxon signed-rank test (a) or the one-sided Student's t-test (d, e); $^{*}p < 0.05$ and $^{**}p < 0.01$. Correlation *r* and its *p*-value were calculated using the Spearman correlation (b).

#### Extended Data Fig. 7. Enhanced sensitivity to anti-mitotic drugs upon DBR1 depletion

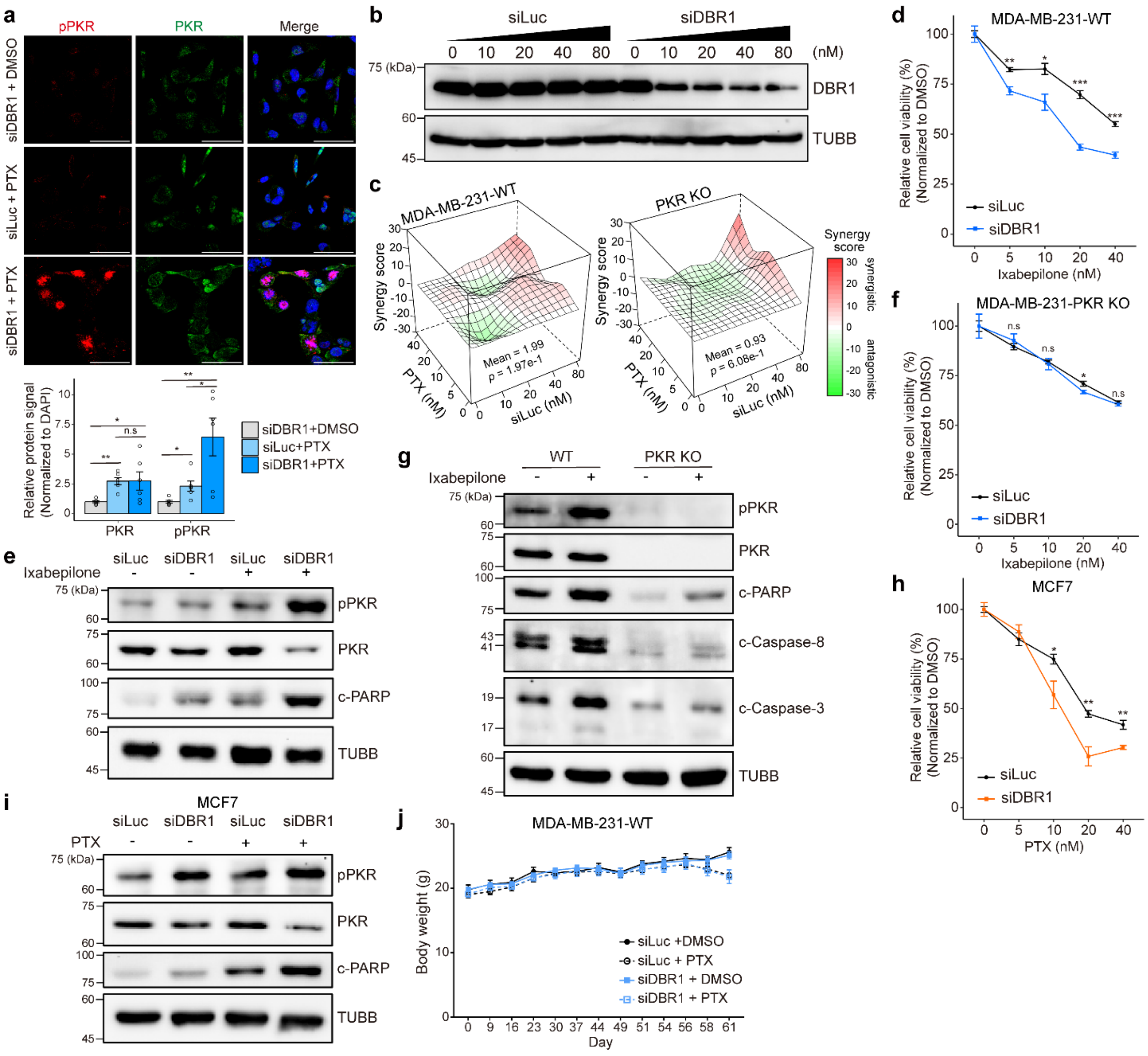


**a.** PKR phosphorylation upon 20 nM PTX treatment in siRNA-transfected MDA-MB-231, analyzed by immunocytochemistry. Quantification is shown below (*n* = 6). Scale bar indicates 50 μm.

**b.** WB of DBR1 knockdown efficiency across increasing siDBR1 concentrations.

**c.** Synergy scores between siLuc and PTX in MDA-MB-231 WT and PKR KO, calculated using the Bliss independence model. Mean synergy scores and p-values are indicated. Positive and negative scores indicate synergistic and antagonistic interactions, respectively.

**d.** Cell viability upon the treatment with various concentrations of ixabepilone (*n* = 4 for each concentration) in siRNA-transfected MDA-MB-231. Cell viabilities are normalized to their respective values at no ixabepilone treatment condition.

**e.** PKR phosphorylation and PARP cleavage upon 20 nM ixabepilone treatment in siRNA-transfected MDA-MB-231, analyzed by WB.

**f.** Cell viability upon the treatment with various concentrations of ixabepilone (*n* = 4 for each concentration) in siRNA-transfected MDA-MB-231 PKR KO. Cell viabilities are normalized to their respective values at no ixabepilone treatment condition.

**g.** WB of apoptotic markers upon 20 nM ixabepilone treatment in siDBR1-transfected MDA-MB-231 WT and PKR KO.

**h.** Cell viability upon the treatment with various concentrations of PTX (*n* = 4 for each concentration) in siRNA-transfected MCF7. Cell viabilities are normalized to their respective values at no PTX treatment condition.

**i.** PKR phosphorylation and PARP cleavage upon 20 nM PTX treatment in siRNA-transfected MCF7, analyzed by WB.

**j.** Body weight curves of NOD/SCID mice (MDA-MB-231 WT xenografts) treated with nanoparticle-formulated siLuc or siDBR1, followed by DMSO or PTX intraperitoneal administration. Body weights were measured at the indicated time points and are presented as mean ± s.e.m. (*n* = 4 per group).

All error bars denote s.e.m. *P*-values were calculated using the one-sided Student's t-test (a, d, f, h); **p* < 0.05, ***p* < 0.01, ****p* < 0.001, and n.s: non-significant.

### Extended Data Fig. 8. Effects of ADAR1-mediated A-to-I editing on intron lariat RNAs

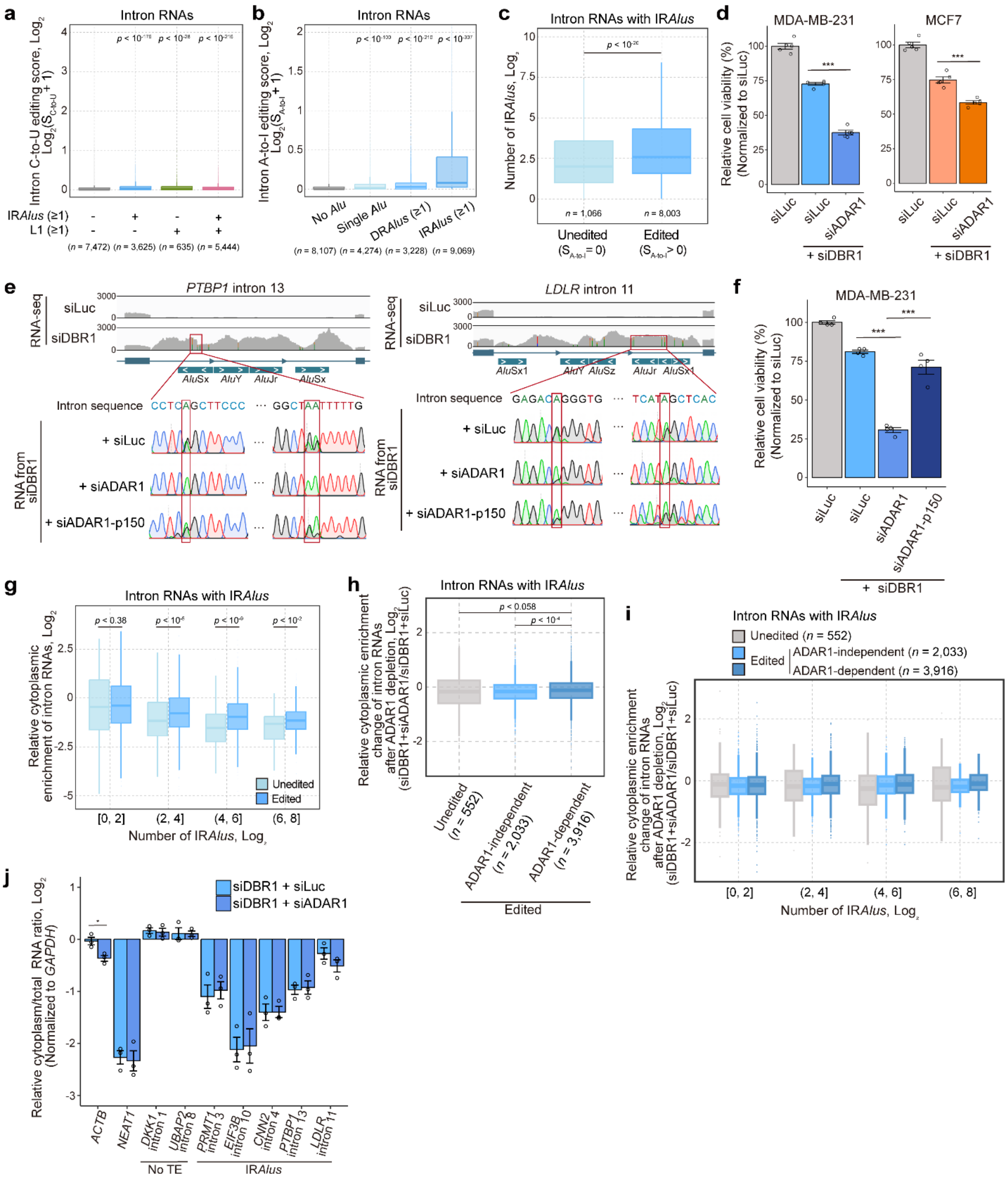

**a.** C-to-U editing scores of intron lariat RNAs depending on the presence of either IR*Alus* or L1.
**b.** A-to-I editing scores of intron lariat RNAs depending on the type of *Alu* associations (Single *Alu*, DR*Alus*, and IR*Alus*). Intron RNAs lacking *Alu* elements are categorized as “No *Alu*”.
**c.** Box plot for the number of IR*Alus* for A-to-I edited intron RNAs (A-to-I editing score > 0) and unedited intron RNAs (A-to-I editing score = 0). The number of IR*Alus* for each intron RNA was calculated by multiplying the counts of sense and antisense *Alu* elements.
**d.** Effect of ADAR1 depletion on cell viability in DBR1-deficient MDA-MB-231 and MCF7 ($n = 5$).
**e.** A-to-I editing rate change of *PTBP1* intron 13 and *LDLR* intron 11 after ADAR1 or ADAR1-p150 depletion in DBR1-deficient MDA-MB-231, confirmed by Sanger sequencing.
**f.** Effect of ADAR1-p150 depletion on cell viability in DBR1-deficient MDA-MB-231 ($n = 5$).
**g.** Comparison of relative cytoplasmic enrichment for A-to-I edited and unedited IR*Alus*-containing intron RNAs, categorized by the number of IR*Alus*.
**h, i.** Effect of ADAR1 depletion on the relative cytoplasmic enrichment for A-to-I edited and unedited IR*Alus*-containing intron RNAs, classified based on ADAR1 dependency of A-to-I editing (**h**) and the number of IR*Alus* (**i**).
**j.** Effect of ADAR1 depletion on the relative cytoplasm/total ratio of intron RNAs analyzed by RT-qPCR following subcellular fractionation in DBR1-deficient MDA-MB-231 ($n = 3$).
All error bars denote s.e.m. *P*-values were calculated using either the two-sided Wilcoxon signed-rank test (a, b, c, g, h) or the one-sided Student’s t-test (d, f, j); $^{*}p < 0.05$ and $^{***}p < 0.001$.

**Table 1. Sequences of siRNAs**

| | **Sense (5′-3′)** | **Antisense (5′-3′)** |
|---|---|---|
| siLuc | CUU ACG CUG AGU ACU UCG A | UCG AAG UAC UCA GCG UAA G |
| siDBR1_1 | CUG AUC AAG CUU CUG AGU U | AAC UCA GAA GCU UGA UCA G |
| siDBR1_2 | AGA GCA UCU CAA ACC UAC U | AGU AGG UUU GAG AUG CUC U |
| siDBR1_3 | CCA UGU AAC UUU AGU GUA A | UUA CAC UAA AGU UAC AUG G |
| siDBR1 (3′ UTR) | GGA CCA UAA AGA UGA GUA A | UUA CUC AUC UUU AUG GUC C |
| siPKR_1 | GCA GGG AGU AGU ACU UAA A | UUU AAG UAC UAC UCC CUG C |
| siPKR_2 | GCA UGG GCC AGA AGG AUU U | AAA UCC UUC UGG CCC AUG C |
| siMDA5 | GAA AAU GCA UCA CGU CAA U | AUU GAC GUG AUG CAU UUU C |
| siRIG-I | CUC UUG AUG CGU CAG UGA U | AUC ACU GAC GCA UCA AGA G |
| siRNase L | GAA CUA CAC UUG UCC UGU G | CAC AGG ACA AGU GUA GUU C |
| siNF90/NF110 | CUG ACA AAG CAC GGC AAG A | UCU UGC CGU GCU UUG UCA G |
| siPACT | AGG AAU GCU GCU GAG AAA | UUU CUC AGC AGC AUU CCU |
| siTRBP | CAC GUC AGC UAC CUG GAU A | UAU CCA GGU AGC UGA CGU G |
| siADAR1 | GAG AAA AGC UCC AAA UAA A | UUU AUU UGG AGC UUU UCU C |
| siADAR1-p150 | GCC UCG CGG GCG CAA UGA A | UUC AUU GCG CCC GCG AGG C |

**Table 2. Sequences of shRNAs**

| | **Sequences (5′-3′)** |
|---|---|
| shScramble | CCU AAG GUU AAG UCG CCC UCG CUC GAG CGA GGG CGA CUU AAC CUU AGG |
| shDBR1 | GCG AGC UGG AUA AGA UCU AUG CUC GAG CAU AGA UCU UAU CCA GCU CGC |

**Table 3. RT-qPCR primer sequences**

| | **Forward (5′-3′)** | **Reverse (5′-3′)** |
|---|---|---|
| *ID1* intron 1 | TATACCGACGGGGAAACGGA | TTTCCAGGCTCCTTAGGCAC |
| *DKK1* intron 1 | GCTGAATGTGTGCGGTTCAG | GGCGCTGATCACAGTCCTTA |
| *UBAP2* intron 8 | CCCTGGTGGTCTTCTTCCATC | CACCCGGGAATGGGATCTG |
| *MYH9* intron 25 | TGTCTCCCCAGCAAAAGCAC | GCGTATCTCCTAAGCGAGCC |
| *PTBP1* intron 13 | GCACCTCTTACCCCAAACCT | TCCAAAGTGTCCAACGACCC |
| *LDLR* intron 11 | TCAGTGTGGTGCTGACAACC | ACACCAAGTGTGCCTTCTCC |
| *ACTN4* intron 16 | AGAGAGCTCCCGTAGTGAGG | AGAATTGGGACCTGCTCTGC |
| *RDH11* intron 4 | ATCCCAGGGCAGAGGTACATA | TTGTTGCATCCACCTCTCTCAA |
| *TARDBP* intron 1 | GAGCGTCGTTAGGTCCTGG | CAGCGTCCCTCGAAGTACAA |
| *MYH9* intron 27 | TTAGCTTGGCTGGTTCCTGG | GAGTGGAGAAAGGAGGCTCG |
| *CCNF* intron 8 | ACCTGAGCCATCCTTGTTCC | CCCATTCAGGCACCAAATGC |
| *PRMT1* intron 3 | CTGCGCACTTCCTAGGGTC | CCCTGCAGAGATTCAGGCTAA |
| *CANX* intron 5 | TTGCCACCTTTGTGTGTTGC | TCCAACATTCAGTAGCACAGGT |
| *DHX9* intron 12 | AGTGGCATGCAGTCTGAGAA | CAGGACAGGCATTTTGTGTGG |
| *PSMC6* intron 6 | ACAGTGATTTGCTGCATTAGCC | AAAGCGTGACTGGGAAAAGC |
| *MYDGF* intron 3 | TCATCCCCTGCTGTCACCTT | GCTTCTCAGAGGGCATTGACT |
| *RNF213* intron 21 | ACTTGGCAGGGTTATGTGAGG | AACCCCCATGGCTGTTTCTC |
| *EIF3B* intron 10 | AACACACAGCTGGAATCGGA | AGGAAGGAGGCCAAGAAACG |
| *TRIO* intron 4 | CCTTCGGCCATTCCTCAGTT | GTGTCCCAGGTCACAGTCAG |
| *UTP18* intron 12 | TATGCGCTGTCCAGGTTCTT | TCCCTAAAGATCCACCTTGTCA |
| *CNN2* intron 4 | GACATTGGAATTGGCTGGGC | CCACGTGAGGTTACTGGGAG |
| *GAPDH* | CTCCTCCACCTTTGACGCTG | TCCTCTTGTGCTCTTGCTGG |
| *ACTB* | CCTGTACGCCAACACAGTGC | ATACTCCTGCTTGCTGATCC |
| *18S rRNA* | CGCCGCTAGAGGTGAAATTCT | CGAACCTCCGACTTTCGTTCT |
| *NEAT1* | GGAGGGCCGGGAGGGCTAAT | CGGTCAGCCCCGTCGAGCTA |
| *RNASEL* (for RNase L) | AAGCATTTCCCCCAAACCCA | ATCAGTCCATCAGCACCCAG |
| *ILF3* (for NF90/NF110) | TGTAAGGAGAAGCCCACAAC | GTCAGGTGGATGGTCAGGGA |
| *PRKRA* (for PACT) | CCCTTAATGCCTGACCCTTCCA | CAGGAAGTCTCCAGCCATGATG |
| *TARBP2* (for TRBP) | AAGACGCCTGTGTACGACCTTC | TTGTGCTTGGCTGCCTTCTTGC |
| *IL6* | CCTTCTCCACAAACATGTAACAAGA | TCACCAGGCAAGTCTCCTCA |

| | | |
|---|---|---|
| *IL8* | ATGACTTCCAAGCTGGCCGTGGCT | TCTCAGCCCTCTTCAAAAACTTCTC |
| *CXCL10* | AGCAGAGGAACCTCCAGTCT | ATGCAGGTACAGCGTACAGT |

**Table 4. Lariat-specific RT-qPCR primer sequences**

| | **Forward (5′-3′)** | **Reverse (5′-3′)** |
|---|---|---|
| *ID1* intron 1 | CCACTTCCGTCCCATCCTT | GGTCGGATCTGGATCTCACT |
| *DKK1* intron 1 | GAGGGAGTAGAACGTGCTGA | GCCCGACCCCTCTCACT |
| *UBAP2* intron 8 | CCGTCTCTGCCCCACTGAGC | GAGTTCCGCCATACACTCACT |

**Table 5. Sequences of CIRC template for in vitro synthesis of lariat-mimic circular intron RNAs**

| | |
|---|---|
| **CIRC template sequence(5′-3′)** | |
| TAATACGACTCACTATAG (T7 promoter) -<br>GGAAATAATTGAGCCTTAAAGAAGAAATTCTTTAAGTGGATGCTCTCAAACTCAGGGAAACCTAAA<br>TCTAGTTATAGACAAGGCAATCCTGAGCCAAGCCGAAGTAGTAATTAGTAAGTTAACAATAGATGA<br>CTTACAACTAATCGGAAGGTGCAGAGACTCGACGGGAGCTACCCTAACGTCAAGACGAGGGTAAA<br>GAGAGAGTCCAATTCTCAAAGCCAATAGGCAGTAGCGAAAGCTGCAAGAGAATG (group Ⅰ Intron) -<br>AA (3’ exon) - Payload (Target intron sequence) - TCT(5’ exon) | |
| **Target intron sequence(5′-3′)** | |
| *DKK1*<br>intron 1 | GTGAGAGGGGTCGGGCACTCAGAGGATGCTCTGACCTTGAAAGGGTCCTATCTGGAGAC<br>GAGGGAGTAGAACGTGCTGAATGTGTGCGGTTCAGGGAGCATTTGGTAACCCTGCATTTG<br>GGAGCAGTGGGCAGTAACAGGTTTTGGAGAGGTGGACAGATAAGGACTGTGATCAGCGC<br>CCGGGTCCAAGAGGGCGGGTACCTGGACGTCTGGGTGCCTCA |
| *CANX*<br>intron 5 | GTGAGAATCCCATTTGGTTTAGATATAATAGCAGATAGAGGTTTGACATGTTGTCTCCATG<br>GCATTTCCTAAGACTAGTTTAAAAGGACATAGTAGGCTTTAAGATTTTTCCAgccaggtgcggtg<br>gctcacgcctgtaatcccagcactttcggaagccaaggcgggcagatcacgaggtcaggagatcgagaccatcctggctaacac<br>agtgaaaccccgtctctactgaaaatacaaaaaattagctgggcgtggcagcgtgcatctgtagtcccagctgctggggaggctga<br>ggcaggagaatggtgtgaacccgggaggccaagcttgcagtgagccgagaacgcgccactgcactccagcctgggtgacaga<br>gcgagactccgtctcaaaaaaaaaaaaaaaaaaaGATTTTTCCATGGAAGTTTTGCCACCTTTGTGTGT<br>TGCCTAAttttttttttttttttttttatttAAGGTTCAGGGGTGCATGTGTAGCCTTATATTTAATAGGATTA<br>GATCTGTTTAATCACCTGTGCTACTGAATGTTGGAAATTTCTAGACAGTAGTATAATACACT<br>ACAGTTTTAAAAAGCCATATCTGAATAATATTTGCCAAATAAAGTGTGtttatttattatttatttactttttg<br>agacggagtctcattcttgccacccaggcttctggaatagagtggcgtgatcttggctcaccgcaacctctgtttcttgggttcaagcgat<br>tctcctgcctcagcctaccgagtagttgggattacaggtgcccatcaccacgcccggctaatttttgtgtttttattagagatggggtttcac<br>catgttggccaggctggtctcgaactcctgaccttgagtgatccgcctgccctcagcctccctaagtgctgggattataggtgtgagcg<br>accacgGCTGCCCCAGTATAATATATTTATGCAACTAGGAGGTGTTTTTTTTGGCATTTAAAG<br>GACCTTCA |
| *PTBP1*<br>intron 13 | GTGAGTGCTGGGCCGGGGGGCTCATGGGGCCGGGGGCGGGCAAGGGCTCTGCTTGGC<br>TGTCCTACCGCGTCGGTGTGTGGACTTCTGGCGTTTCCAGAGTGCAGGTCGGGCACCTC<br>TTACCCCAAACCTGAAGTACTGCAAGGCCTGGAGCTTGAGCCGAGGGCTGCTCAAGGGA<br>GGGGGTCGTTGGACACTTTGGAGGTTTTGGCTCAGGGGATGCTCCCTGTGAGAATGTATA<br>ATGCACACAtttatttatttatatttttttgaggcgaagtcttgctttgtcacccaggctggagtacagtggaacgatctcggttcactgc<br>aacctccgcctcttgggttcaagtgattgtgccacctcagcttcccgagtagctgggactgtaggcacccacaaccacacctggctaa<br>tttttgtatttttagtagagacggggtttcaccatattggccaggctgatcttgatctcctgaccttgtgatccgcctgcttcggcctcccaaa<br>gtgctgggatacaggcgtgagccaccgcgcctagccACATTTAtttatttatttatttatttttacttattttgagatggagtcacgctctgt<br>cgcccaggctggagtgcagtggcgggatctctgcaagttctgcctcctgggttcatgccactctgctgcctcagcctcccgagtagctg<br>ggagggactataggtgcccgccagcacgcccggctaatttttttgtatttttagtggaaacggggtttcactgtgttagccaggatggtgtc<br>gatctcctgactttgtgatcaacccgcctcggcctcccaaagtgctgggataacaggtgggagccaccacgcctggccTAATgca<br>cacctttataatcccagtactgtgggggggccgggcaggggattgcttgaacccaagagttaagaccagcctgagcaataaagtgag<br>acctctgtctctacaggaacaattggttaaaattagctaggagtggtggtgtgcacgtcaggtcccagctcttccggattgggaggttga<br>ggtgggaggttcacttcgcgctcaggaggttgaggttgcagtgagctgtgattgcactgccagactccaggctgggcggcagagtaa<br>gaccctgtctgtaaaaaTTACATAATGCAAGTATTTGGAAATGCAGGTACATCTGGCCCCAAGCAT<br>GTCAACCAGTAGTAAGCCCTTTGATATTTAACACGTTACTTGTGAAATGTTATCAGAAAATC<br>AACAAGGggcctggcagagtggctcacgcctatcatcccagcactttaggaggccgaggcaggcagatcacctgaggtcagg<br>agttcaagaccaacctggccaacctggtgaaaccccgtctctactaaaaataaaaaaattagcctggcatggtgttgcgtgcctgta<br>atcccagctacttgggagactgaggcaggagaatcacttgaacctgcaggcggaggttgtggtgagctgagatcacgccactgtgc<br>tcccgcctgggcgacagagcaagattccatctggaaaaaaaaaaGTGGAGGAGCGTGGTACTCTGTTTCTGC<br>AAACTTGTAGATGCCTGGTTTCGTTAGAAGCTGCTTCAGACCATTGTGGACATGATTTGAT<br>ACCCCATTTCTGGGCTCCTAAAAGGCCCTACGCCTTCCCCGGCTACTCTGAAAACTAGTC<br>TGGGGAAAGCCTCGCGGACCTGACTGGGCGCCCCCACCCCCACGCGGCCCCAGGCTCA |
